# A Bacterial Living Therapeutics with Engineered Protein Secretion Circuits To Eliminate Breast Cancer Cells

**DOI:** 10.1101/2023.04.27.538589

**Authors:** Gozeel Binte Shahid, Recep Erdem Ahan, Julian Ostaku, Urartu Ozgur Safak Seker

## Abstract

Cancer therapy can be limited by potential side effects, and bacteria-based living cancer therapeutics have gained scientific interest in recent years. However, the full potential of bacteria as therapeutics has yet to be explored due to engineering challenges. n this study, we present a bacterial device designed to specifically target and eliminate breast cancer cells. We have engineered *Escherichia coli* (*E. coli*) to secrete a Shiga toxin, HlyE, which is a pore-forming protein that binds to HER2 receptors on breast cancer cells. This binding is facilitated by a nanobody expressed on the bacteria’s surface via the Ag43 autotransporter protein system. Our findings demonstrate that the nanobody efficiently binds to HER2+ cells in vitro, and we have utilized the YebF secretion system to secrete HlyE and kill the target cancer cells. Overall, our results highlight the potential of our engineered bacteria as an innovative strategy for breast cancer treatment.

## Introduction

Recent scientific advances have sparked growing interest in the therapeutic potential of bacteria, prompting a paradigm shift in the approach to disease management. With a deeper understanding of the intricate interplay between bacteria and human physiology, the field of bacteria-based therapeutics has gained significant momentum in recent years, presenting new opportunities for innovative treatments^1-3^. Traditionally, bacteria were primarily studied for their pathogenic properties; however, recent advancements in biology and synthetic biology techniques have uncovered their potential therapeutic applications. ^4-6^. The versatility of bacteria has allowed researchers to harness their ability in many ways such as in production and delivery of therapeutic agents to specific tissues ^3,7^. Bacterial therapy offers a significant advantage over traditional drug therapy by allowing bacteria to target diseased tissues specifically ^8^ and cost-effectively mass-produced. ^9^

Traditional cancer treatments, such as chemotherapy and radiation therapy, often come with a range of undesirable side effects including damage to healthy tissue and lack of specificity ^10^. Bacteria, with their natural ability to selectively colonize and proliferate in specific tissues ^11^, have the potential to offer a new way to overcome this challenge ^1,12^. Bacterial vectors designed to deliver cancer-killing agents directly to the site of cancer allow for a more targeted and effective delivery, while reducing the risk of side effects ^1^. Certain bacteria have also been shown to produce compounds that can directly kill cancer cells, or stimulate the body’s immune system to attack the cancer ^13^. Cancer and bacteria immunotherapy have a long and intertwined history, dating back to the 19th century Coley’s experiments ^14^ which led to the observation that certain bacterial infections could lead to remission in cancer patients by inducing an immune response ^15^. Despite their long history, the use of bacteria as therapeutics has yet to be fully explored, and they have not been as widely adopted as traditional drug therapies. This is partly due to the difficulty in engineering bacteria, but with advances in synthetic biology, this challenge can be overcome ^4,6,9,16,17^. The ability to precisely manipulate and control the genetic information of bacteria owing to synthetic biology has opened novel avenues to create tailored anti-cancer therapeutic strategies with improved specificity and efficacy ^18^.

So far synthetic biology has enabled researchers to manipulate bacteria for the purpose of targeting tumor cells with greater precision ^19^. By modifying bacteria to express specific molecules, they can be made to bind to tumor cells ^12^ and introduce payloads such as macromolecules into the cells ^20,21^. Enhancing the delivery efficiency through the addition of certain genes can improve the replication of the bacteria within the cells ^22^. Additionally, bacteria have been engineered to deliver drugs by opening pores in cell membranes or by lysing the cells ^23^ to release drugs within tumor cells ^24^. Other bacteria have been designed to metabolize waste generated by tumors ^25^ and still others have been programmed with a quorum-sensing system to trigger drug release ^26,27^. Scientists have also developed genetic circuits in bacteria to act as controlled delivery systems for releasing cargo to tumor cells ^22,28^. Further streamlining of the genetic circuits has been tested, for example by expression of the invasin gene, which has been shown to promote bacterial uptake into phagosomes and increase the invasion efficiency of otherwise extracellular bacteria ^29^. Despite the progress in developing bacterial therapies, a major challenge remains the lack of *in vitro* tools for testing bacteria in a tumor-like environment. However, recent advancements have enabled the high-throughput and long-term growth of bacteria in the core of tumor spheroids *in vitro* ^30^.

In this study, we present a design for engineered bacteria that addresses two main constraints in cancer therapeutics: attachment and efficient delivery of therapeutic agents. We have engineered *E. coli BL21* BL21 to secrete a toxin and bind to HER2 receptors on breast cancer cells, facilitated by a nanobody ^18^ expressed on the surface of the bacteria via the Ag43 autotransporter protein system ^31,32^. We used anti-HER 2Rs15d nanobody which has been shown to efficiently bind to HER2+ cells both *in vivo* and *in vitro* ^33^. The pore-forming toxin HlyE was secreted using the YebF secretion system, a type VI secretion system characterized by the ability to secrete a wide range of different compounds, including proteins, toxins, and other molecules ^34^.

To determine the efficacy of our proposed design, we tested the effects of our engineered bacteria on JIMT1 HER2+ cells. To enhance accuracy of our hypothesis, we employed cell spheroids which have a number of advantages, including the ability to mimic the complex microenvironments that are present *in vivo*, improved cell-cell interactions, and increased cell viability. In our experiments, we demonstrated that our engineered bacteria are functional and show promising results *in vitro* in both 2D and 3D assay experiments. Our results demonstrate the potential of our approach for engineering bacteria as a novel treatment strategy for breast cancer, and we believe that further experimentation, including *in vivo* studies, will provide even more insight into the potential of this approach.

## Results and Discussion

### Design and Characterization of Engineered Bacteria

We engineered a bacterial device that counters two main problems of cancer therapy: specificity and targeted delivery of therapeutic agent to cancer cells. For our experiments we used the Bl21 strain of *E. coli BL21*, due to the ease of engineering it. To enable specific binding of the bacteria, we used a nanobody (Nb) that targets HER2+ receptors on breast cancer cells.

The binding affinity and specificity of 2Rs15d nanobody to HER2 receptors, both *in vivo* and *in vitro*, has been extensively studied ^33^. To aid attachment of *E. coli BL21* to receptors on mammalian cells, the Nb would have to be displayed on the surface of the bacteria (Figure 1A). We achieved this by using the Ag43 autotransporter protein system which has been shown to successfully display proteins on bacterial surfaces, using a signal peptide ^31,32^.

**Figure 1.**
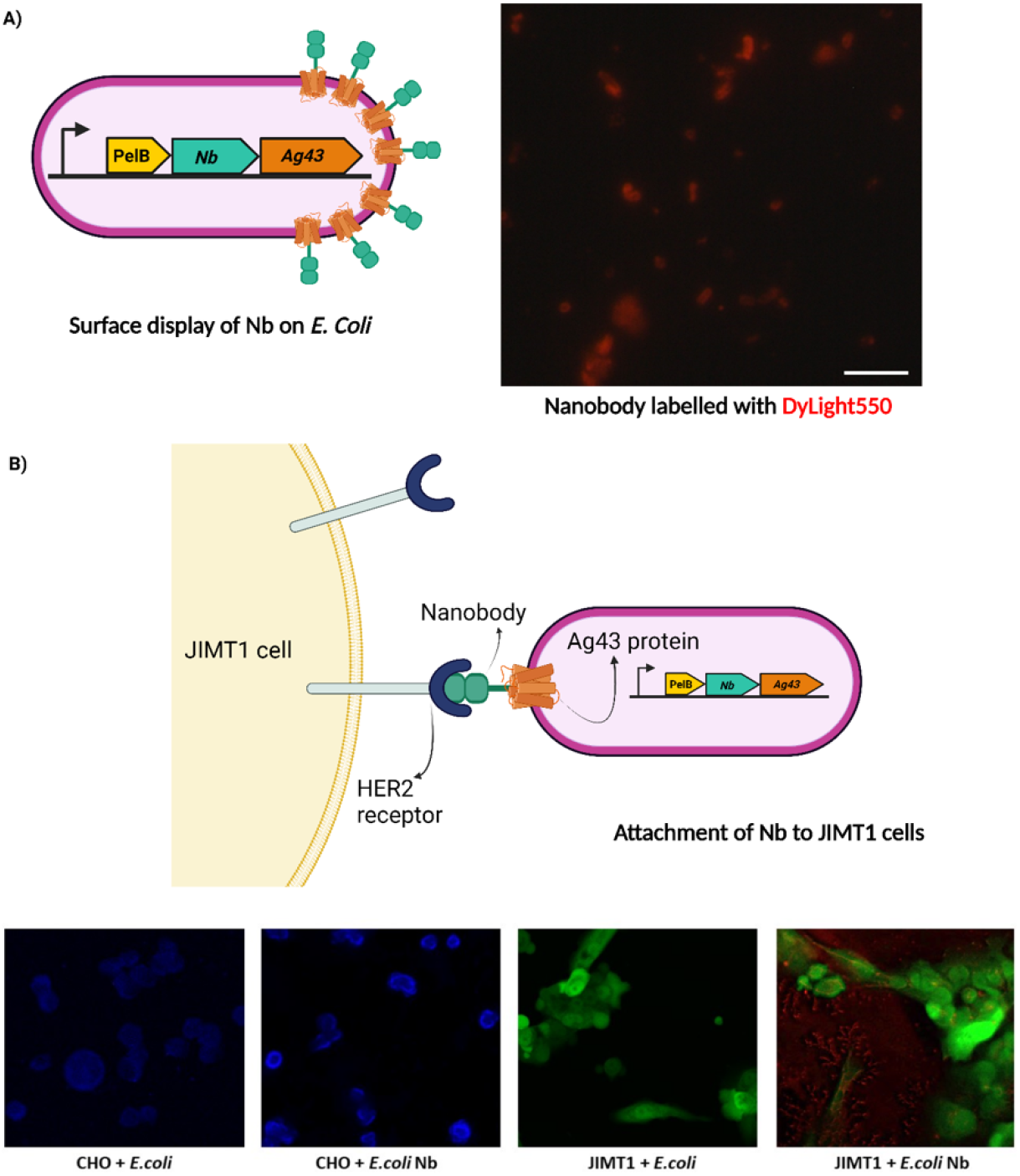
Engineering *E. coli* with surface displayed Nanobody (Nb) *2Rs15d* for binding to JIMT1 breast cancer cells. (A) Bacteria design for surface display of Nb using autotransporter protein *Ag43* system and PelB signal peptide. Immunocytochemistry image showing Nb on bacteria tagged with DyLight 550, giving red fluorescence. (B) Diagrammatic illustration made using Biorender.com to show binding of surface-displayed Nb on *E*.*coli* t HER2 receptors on JIMT1 breast cancer cells. Binding assay was done to verify attachment of *E. coli* Nb to JIMT cells. *E*.*coli* was transformed with pZA mproD RFP for red fluorescence. HER2-CHO cells, used as a control, show no binding of *E. coli* or *E. coli BL21* Nb bacteria. *E. coli BL21* does not show binding with JIMT1 cells, however *E. coli BL21* Nb can be seen (red) bound to JIMT1 cells in the fourth group, due to the presence of Nb.

To verify our engineered *E. coli BL21 design* with surface-displayed Nb, we performed a Western Blot using whole cell lysate and got the relevant band at ∼40 kDa (Supplementary Figure 1). To confirm the presence of the Nb on the bacterial surface, we conducted an immunocytochemistry experiment using 2Rs15d tagged with DyLight 550, which produces red fluorescence. Images were taken with inverted light microscope; the presence of red fluorescence indicates presence of Nb on bacteria. The Immunocytochemistry image in Figure 1A shows successful display of nanobody on the surface of our engineered *E. coli BL21*.

We hypothesize these bacterial cells to bind to JIMT1 mammalian breast cancer cells (Figure 1B) due to overexpression of HER2 receptors in JIMT1. To test this we performed a binding assay using CHO-K1 cells as a negative control. *E. coli BL21* and *E. coli BL21* with surface-displayed nanobody (*E. coli BL21* Nb) bacteria were incubated with both CHO and JIMT1 cells seeded in a well-plate with glass slides at the bottom. After one hour of incubation the slides were washed with PBS to remove any unbound bacteria (Supplementary Figure 2). Confocal microscopy images in Figure 1B show CHO cells (blue) with no bound *E. coli BL21* or *E. coli* Nb. JIMT1 cells (green) show no binding of *E. coli*, however *E. coli* Nb can be seen (red) bound to JIMT1 cells, confirming our hypothesis.

**Figure 2.**
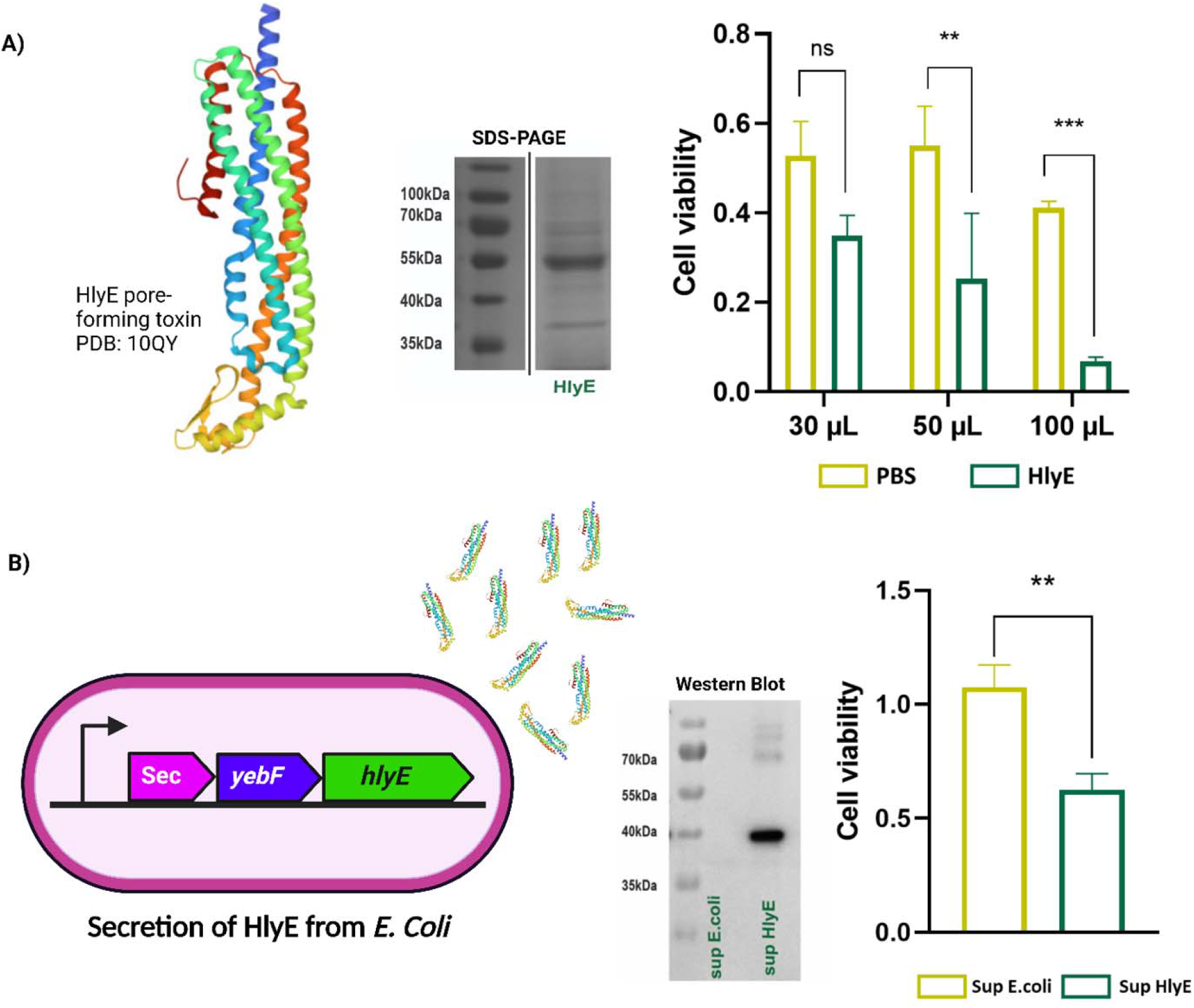
Pore-forming HlyE protein as an anti-cancer agent. (A) X-ray crystal helical bundle structure of HlyE (PDB:10QY). SDS-Page to show purified HlyE protein band at 55kDa. Purified HlyE protein and PBS (control) is added to JIMT1 mammalian cells in varying volumes to assess killing of cells by HlyE. Cytotoxicity assessment of purified HlyE is done by MTT (3-[4,5-dimethyl-2-thiazolyl]-2,5-diphenyl-2H-tetrazolium bromide) assay. (B) *E. coli BL21* design for secretion of HyE using *YebF* secretion system. Western Blot shown for verification of secretion (HlyE band [40kDa) is done by using supernatant samples concentrated by TCA precipitation. Supernatant of *E. coli BL21* secreting HlyE is added to JIMT1 cells to assess its toxicity, and supernatant of *E. coli BL21* is added as a control. Cytotoxicity assessment of HlyE secreted by engineered *E. coli BL21* is done using MTT (3-[4,5-dimethyl-2-thiazolyl]-2,5-diphenyl-2H-tetrazolium bromide) assay.

For the therapeutic agent, we decided to use pore-forming Hemolysin E (HlyE) which is known to exhibit cytotoxic effects in mammalian cells ^35^. We first purified the HlyE protein to test its validity as a toxic agent for JIMT1 cells. Purification verification was done by SDS-PAGE and we got the relevant band at ∼55 kDa (Figure 2A). To determine the effect of purified HlyE on our JIMT1 cell line we added various volumes (30 µl, 50 µl, and 100 µl) to mammalian cells seeded in a well-plate. PBS was added to half the wells in equal volumes as a negative control. After incubating the well plates we performed MTT assay for testing cell viability; our graph (Figure 2A) shows JIMT1 cells treated with HlyE were significantly lowered as compared to those treated with PBS, especially at higher volumes. These results show our choice of HlyE toxin to be a viable therapeutic agent for breast cancer cells. We then engineered our *E. coli BL21* to secrete HlyE using YebF secretion system (Figure 2B). The YebF secretion system belongs to the type VI secretion system category, known for its diverse secretion capacity of various compounds ^34^. After cloning and sequencing our bacteria, we did a Western Blot to test its supernatant for presence of HlyE using TCA precipitation method, and got the relevant band at ∼40 kDa (Figure 2B). To test the efficacy of our HlyE secreting *E. coli BL21* we incubated JIMT1 cells with the supernatant of both *E. coli BL21* and *E. coli BL21* HlyE and performed a cell viability assay using MTT. Our MTT results in Figure 2A show a significant decrease in JIMT1 cells incubated with *E. coli BL21* HlyE supernatant as compared to *E. coli BL21* alone. The fundamental objective of this research endeavor was to employ genetic engineering techniques to create a novel bacterial construct that effectively amalgamates the surface-expressed Nb and the HlyE secretion pathway (Figure 3A). This integrated system is envisioned to offer synergistic benefits, where the specific binding ability of the Nb can be leveraged to selectively target and bind to JIMT1 cells, and subsequently, the secretion of HlyE can facilitate the killing of target cells.

**Figure 3.**
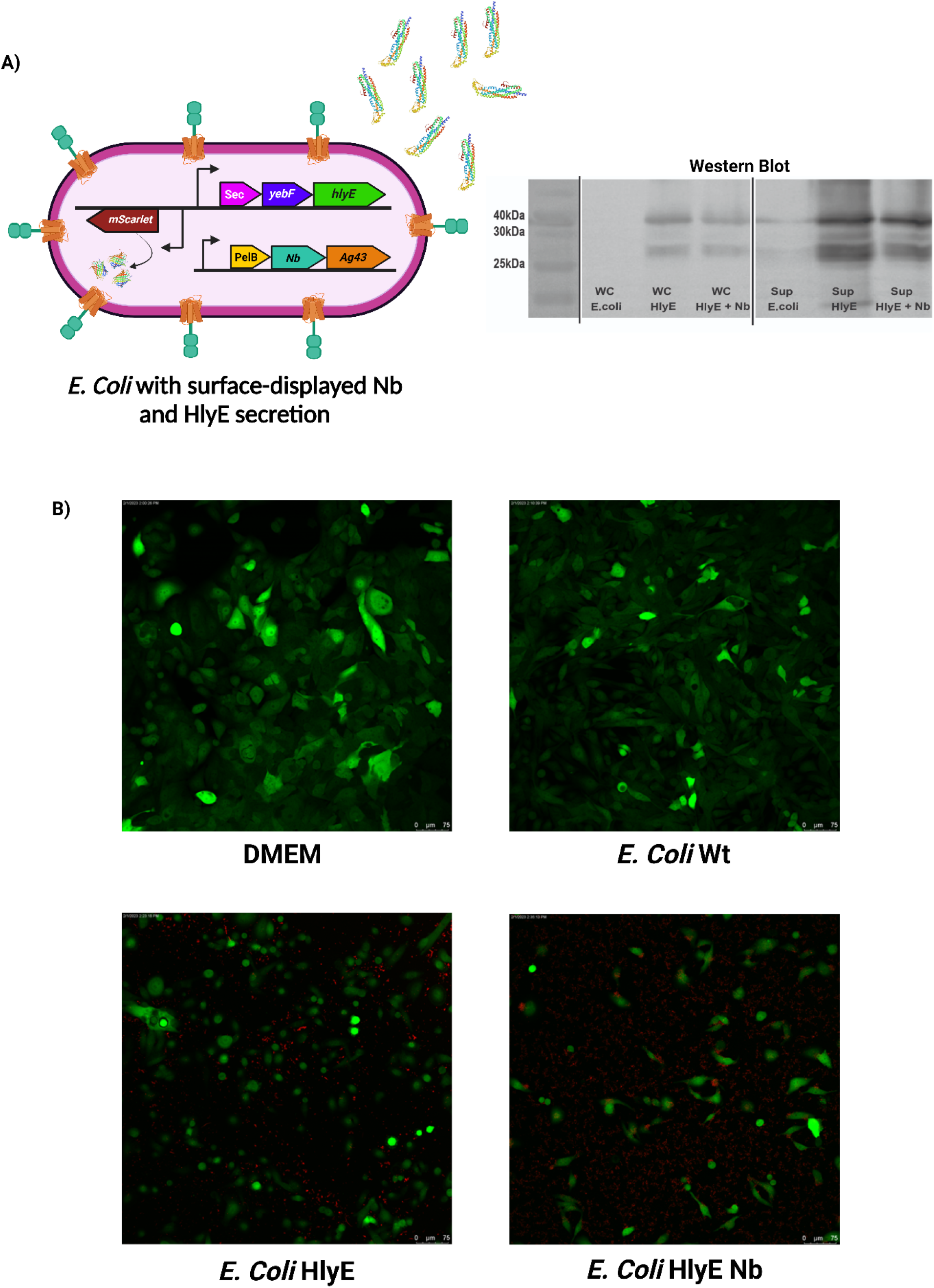
*E. coli BL21* with surface display of Nb and HlyE secretion tested on JIMT1 mammalian cells. (A) Bacteria design for surface display of Nb and secretion of HlyE toxin. Western Blot with whole cell (WC) and supernatant (Sup) concentrated with TCA precipitation showing bands at 40kDa for HlyE in all engineered bacteria cells. (B) 2-D assay experiment with green fluorescent JIMT1 cells and red fluorescent *E. coli BL21* cells. Images show JIMT1 cells in normal conditions DMEM, one day after incubation with *E. coli BL21* (control), and after incubation with *E. coli BL21* HlyE and *E. coli BL21* HlyE Nb. Killing of JIMT1 cells is clearly observed in experimental groups.

### Examining the Impact of Genetically Engineered Bacteria on JIMT1 Breast Cancer Cells

After optimizing our engineered bacteria design, we aimed to comprehensively characterize the impact of our genetically engineered bacterial construct on the cellular behavior and viability of JIMT1 breast cancer cells. We verified that our engineered *E. coli BL21* expresses the desired proteins by performing a Western Blotting experiment with both Whole Cells (WC) and supernatants (sup), using unaltered *E. coli BL21* as a control (Figure 3A). The addition of mScarlet to bacteria was done with the aim of enhancing contrast and facilitating visualization during imaging experiments. To ascertain *in vivo* effects of the bacteria we decided to perform a 2D cell culture experiment. JIMT1 cells cultured with standard growth medium DMEM and with *E. coli BL21* Wild-type (Bl21) were used as a negative control. In order to test the efficacy of our double-transformed bacteria we included *E. coli BL21* secreting HlyE group to determine whether *E. coli BL21* HlyE Nb indeed performs a more targeted release of HlyE than *E. coli BL21* HlyE resulting in greater diminution of JIMT1 cells. Mammalian cells seeded in a 96-well plate were incubated overnight with relevant bacteria and media added. Confocal microscopy images were taken the next day showing a negligible difference in mammalian cells in *E. coli BL21* Wt group as compared to those growing in DMEM, a more significant difference in *E. coli BL21* HlyE group and the most difference in terms of quantity and morphology of JIMT1 in *E. coli BL21* HlyE Nb group (Figure 3B). Our JIMT1 cells are stably transfected with GFP and can be seen in green, whereas the bacteria with mScarlet can be seen in red. We also performed MTT assay after killing bacteria in the wells with Gentamycin. To quantify cell viability results we performed a one-way Annova, with *E. coli BL21* Wt as the control group (Supplementary Figure 3) The results show that difference between *E. coli BL21* Wt and *E. coli BL21* HlyE is non-significant, whereas the difference between *E. coli BL21* Wt and *E. coli BL21* HlyE Nb is substantial. Based on the findings obtained from this experimental study, it can be inferred that our genetically modified bacterial strain has demonstrated an effective cytotoxic activity against JIMT1 breast cancer cells.

After careful analysis, we determined that relying solely on a single assay was insufficient in determining the efficacy of our system. To better replicate the complex environment of a tumor cell mass and bolster the validity of our hypothesis, we opted to conduct 3D cell culture experiments by creating JIMT1 cell spheroids. Spheroids grown in 3D cell culture represent a sophisticated and intricate cell culture model that is characterized by its increased physiological relevance and complexity compared to conventional 2D cultures. In contrast to 2D monolayers, 3D cell culture spheroids are composed of clusters of cells that are able to establish intricate cell-cell and cell-matrix interactions resulting in enhanced cell differentiation, proliferation, and survival, which better resemble the *in vivo* microenvironment of a tumor mass ^36,37^. Therefore cell spheroids provided a superior and more biologically relevant tool for our study. JIMT1 cell spheroids were prepared using high-purity agarose method ^38^, whereby the bottom of the wells in a well-plate are first filled with high-purity agarose to create a sort of concaved hydrogel. When cells are added to this, they tend to cluster together at the center, and due to cell-cell plus cell-matrix interactions form a spheroid (Supporting Figure 4A). We formed spheroids ranging between 300-500 µm in diameter over a three-day incubation period. Similar to the 2D assay experiment we had four groups: DMEM, *E. coli BL21* Wild-type (Wt), *E. coli BL21* HlyE, and *E. coli BL21* HlyE Nb (Supporting Figure 4B). Images were taken using confocal microscopy before adding bacteria to the wells once the spheroids were formed, for later comparison to starting state of spheroid. Following this, bacteria was added to the wells; images were taken again four hours after incubation with bacteria (on day 1), the following day (day 2), and a final set of images was taken on day 3. The progression of spread of bacteria and subsequent decrease in spheroid area can be seen in Figure 4. The overlay confocal images show JIMT1 cells in green, bacteria in red, and the black spots indicate dead JIMT1 cells. We noticed a slight decrease in spheroid size in wells that contained DMEM alone with no bacteria. Similarly, spheroids treated with non-engineered *E. coli BL21* showed a meagre gradual decrease over the experimental time period. In comparison, spheroids treated with *E. coli BL21* HlyE showed a much more significant change; the spheroids had lost their cluster formation by day 3, and evident spheroid disruption was visible with much more decreased amount of green JIMT1 cells as compared to the start of the experiment. Changes had becomes visible by day 2 in this case. The largest amount of decrease in spheroid area as well as disruption of JIMT1 cells from a spheroidal shape to scattered cells was observed in JIMT1 group treated with *E. coli BL21* HlyE Nb over the three-day period. We had already tested HlyE as a potent toxin for killing mammalian cells, however our 3D cell culture experiment revealed that the surface-displayed nanobody allows a more targeted release of the toxin, thereby eliminating most of the *in vitro* spheroid mass. We expect that such a system could allow the therapeutic agent to be effectively delivered to tumor cells due to specific binding, while minimizing or subduing the effects of the agent on healthy tissue.

**Figure 4.**
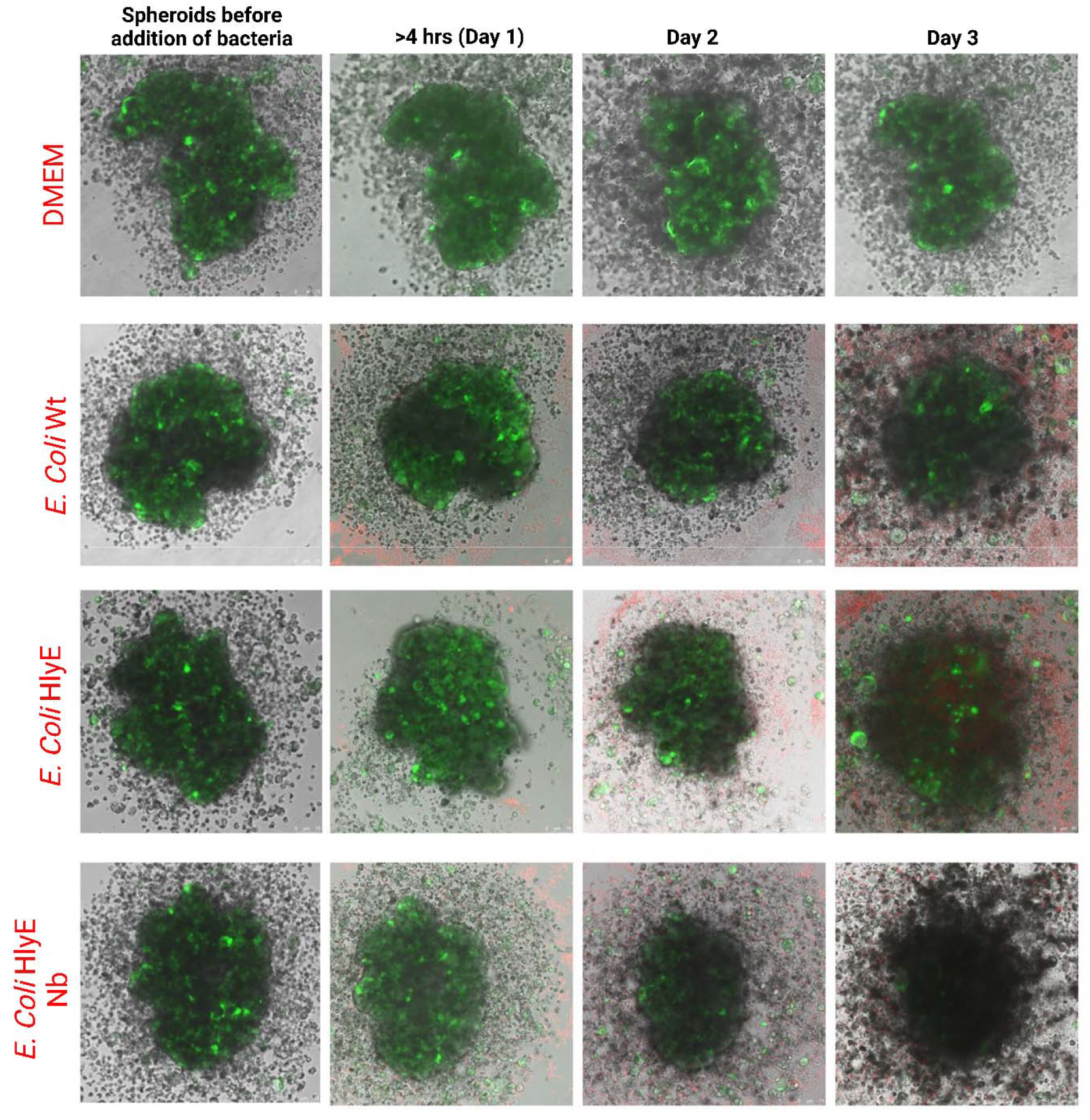
3D cell culture with JIMT1 cell spheroids. Images are taken before addition of bacteria, 4 hours after addition of bacteria (Day 1), on Day 2, and on Day 3.

### Analysis of Data from 3D Cell Culture Experiments

To quantify the data obtained from our spheroid experiment, we conducted statistical computations. We analyzed a total of four replicates of images from each day and group using ImageJ software. In brief, the images were first adjusted for color threshold using the LAB setting after defining a global scale. The spheroid area was then measured using the ‘Analyze Particles’ option. The threshold was adjusted in a standardized manner to eliminate noise depending on image exposures. In cases where there were scattered cells on Day 3 in *E. coli BL21* HlyE and *E. coli BL21* HlyE Nb groups, the ‘Despeckle’ setting was adjusted for to ensure accuracy. After noting the areas for each day, the results were normalized by calculating the % area compared to the starting spheroid area. The normalized results, presented in Figure 5A, indicate a clear trend both among days and groups. The graph illustrates that four hours after induction with bacteria, the spheroid sizes remained relatively stable. However, on Day 2, a more pronounced reduction in spheroid area was observed, with a substantial decline observed in both *E. coli BL21* HlyE and *E. coli BL21* HlyE Nb groups. Furthermore, as evident from the images, even control DMEM conditions showed a progressive decline in spheroid area by Day 3, and the *E. coli BL21* Wt group followed a trend akin to the DMEM group. A further and more evident decrease in spheroid area was observed in the *E. coli BL21* HlyE and *E. coli BL21* HlyE Nb groups on Day 3. The most significant decrease was observed in JIMT1 spheroids treated with *E. coli BL21* HlyE Nb on all four days. Taken together, these findings strongly suggest that the use of our proposed genetically engineered bacteria with nanobody and toxin resulted in the most significant impact on the spheroids of JIMT1 cells compared to the other groups.

**Figure 5.**
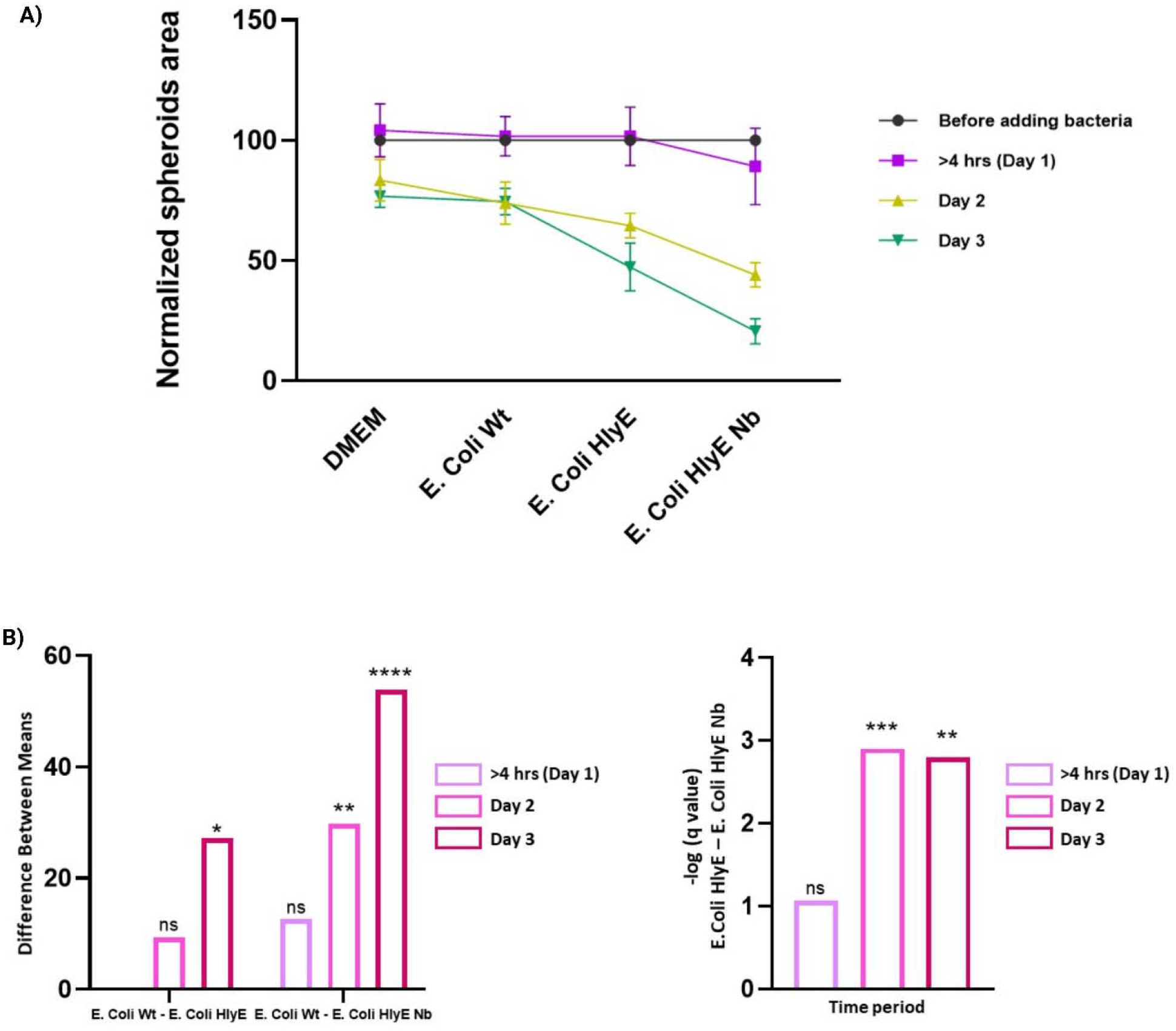
Statistical analyses of spheroids experiment. (A) Image J analysis of JIMT1 spheroids over the course of the experiment showing decrease in normalized spheroids area over a three-day period. (B) 2-way ANNOVA (Dunnett’s Muliple Comparisons Test, p < 0.05 = ns, p < 0.0022 = **, p < 0.0001 = ****, p < 0.0001 = ***) was done with *E. coli BL21* Wild-type (WT) as control group. Multiple t-tests statistical analysis is done to compare cell viability of *E. coli BL21* HlyE group with *E. coli BL21* HlyE Nb group over the three-day peroid (p < 0.05 = ns, < 0.005 = **, p < 0.0005 = ***).

To expand on our results we decided to perform further statistical tests to determine the significance of difference in normalized spheroids area between groups. We performed a two-way Annova using Dunnett’s Muliple Comparisons Test (p < 0.05 = ns, p < 0.0022 = **, p < 0.0001 = ****, p < 0.0001 = ***), with DMEM as the control group (Supplementary Figure 5). The difference between DMEM and all groups was non-significant 4 hours after incubation, with difference between area in DMEM and *E. coli BL21* Wt non-significant over all three days. On day 2 and day 3 *E. coli BL21* HlyE showed a significant decrease in spheroids area (*) as compared to DMEM. *E. coli BL21* HlyE Nb also showed a significant difference on day 2 (*) and a highly significant difference on day 3 (****). Figure 5B shows the same two-way Annova done with *E. coli BL21* Wt as the control group. The difference between *E. coli BL21* Wt and *E. coli BL21* HlyE after 4 hours is negligible, a non-significant difference is reported between these two groups on day 2, but day 3 shows a significant difference (*). *E. coli BL21* HlyE Nb being our main experimental group, as expected shows a larger significant difference as compared to *E. coli BL21* Wt on day 2 (**) and day 3 (****). These Annova results further consolidate our earlier findings and observations. In addition, we also decided to compare the two main experimental groups, *E. coli BL21* HlyE and *E. coli BL21* HlyE Nb, with multiple t-tests (p < 0.05 = ns, p < 0.005 = **, p < 0.0005 = ***) in order to account for differences over all three days. On day two we saw the greatest difference between decrease in spheroids area amongst the two groups (***). However, day 3 results also report a statistically significant difference (**), which reinforces our hypothesis and the potency of our bacteria design.

### Comparative Analysis of the Benefits and Limitations of Our System

Cancer therapy is plagued with a multitude of side effects that often reduce the quality of life of patients undergoing treatment ^10^. To address this issue, we sought to develop a novel alternative or adjuvant to traditional cancer therapies by creating engineered bacteria that can specifically target cancer cells and deliver a therapeutic agent directly to the tumor site, while minimizing collateral damage to healthy tissue. Our approach aimed to overcome the major hurdles in cancer therapeutics of achieving specificity and targeted delivery of therapeutic agents. To this end, we demonstrated the successful display of a HER2 binding nanobody, 2Rs15d, on the surface of our engineered bacteria using the Ag43 autotransporter protein. In addition, we verified the cytotoxic effects of HlyE toxin on a model breast cancer cell line, JIMT1. We further engineered bacteria that display the nanobody and secrete HlyE using the YebF secretion system. Our engineered bacteria were tested on both 2D and 3D cell culture models, and we obtained promising results, highlighting the potential of this approach as an innovative cancer therapeutic.

Unlike traditional chemotherapy drugs that can trigger drug resistance mechanisms in cancer cells, our bacteria-based approach can overcome this problem by directly targeting and killing cancer cells through a different mechanism. There is a growing interest in exploring such bacteria-based therapies that can precisely target cancer cells while leaving healthy tissues unharmed. In this regard, researchers have been developing bacteria that can release therapeutic agents at the site of the tumor using quorum sensing ^26,27^ or enhance their replication in cancer cells by streamlining their genetic circuits ^22^. For instance, the expression of the *inv* gene has been shown to promote bacterial uptake into phagosomes, thus increasing the efficiency of extracellular bacteria invasion ^29^. Building upon these established bacteria systems, we propose a novel and straightforward approach based on nanobody binding and subsequent release of toxin. However, we recognize that some *in vitro* studies do not necessarily translate into *in vivo* results. To address this, we utilized a 3D cell culture model of breast cancer cell spheroids, which more closely resembles the *in vivo* tumor microenvironment. Our choice of *E. coli BL21* Bl21 was motivated by its ease of engineering and replication, as well as its widespread use as a standard laboratory strain. A common laboratory strain of *E. coli BL21* can also easily be scaled up and produced in large quantities, making it potentially more affordable than other therapies. Notably, our approach is also adaptable to other cancer types, with nanobodies and toxin payloads being customized for specific cancer cells. This flexibility allows for a broad range of potential therapeutic applications.

To enhance the comprehensiveness of our research, *in vivo* experimentation is warranted to evaluate the system’s efficacy in the presence of the host’s immune response and other physiological and biological factors, which may exert confounding effects. In addition, genetic circuits with additional elements, such as the *inv* gene, may be incorporated to facilitate complete tumor invasion and colonization by the bacterial strain. While further genetic engineering interventions may pose a potential drawback to the bacteria’s replicative ability, an optimization strategy can be implemented to yield more favorable outcomes. Moreover, *Salmonella*, a bacterium with desirable features as an anti-tumor agent ^39^, can be engineered utilizing our system, since it has demonstrated the ability to proliferate within tumors and elicit tumor regression. Furthermore, while our nanobody is specifically designed to bind to cancer cells and selectively release the HlyE toxin, it is imperative to investigate any potential off-target effects of the toxin on non-cancerous cells. The introduction of bacteria into a live organism may engender a complex and dynamic response, thereby necessitating a comprehensive assessment of the system’s biological outcomes *in vivo*.

## Conclusion

Recent preclinical studies have shown that bacteria-based therapeutics are an effective approach for treating drug-resistant tumors, which are notoriously challenging to treat with traditional therapies. By selectively targeting cancer cells while sparing healthy ones, bacteria-based therapies have the potential to significantly reduce the side effects commonly associated with standard cancer treatments. Moreover, bacteria-based therapies can penetrate deep into tumor tissue, which is frequently inaccessible with conventional methods, providing a key advantage. Although obstacles still exist in developing bacteria-based therapies for clinical use, cutting edge research is being done in this regard for the future of cancer treatment. With ongoing advancements in genetic engineering and biotechnology, researchers are developing increasingly sophisticated bacteria-based therapies that offer even greater specificity and potency in targeting cancer cells. For instance, bacteria can be engineered to produce a variety of therapeutic agents, such as cytokines and checkpoint inhibitors, which can enhance the body’s own immune response against cancer. Furthermore, researchers are exploring the use of bacteria-based therapies in conjunction with other treatment modalities, such as radiation and immunotherapy, to improve their efficacy in treating cancer. Bacteria-based therapies are also being investigated as a potential method for delivering targeted therapies directly to the brain, where conventional therapies have limited effectiveness due to the blood-brain barrier.

In this research endeavor, we have explored the use of bacteria-based therapeutics in conjunction with nanotechnology, which presents the opportunity for the precise delivery of therapeutic agents to specific cells and tissues. To this end, we utilized a biological nanobody (Nb) to facilitate the attachment of *E. coli* to JIMT1 breast cancer cells. Furthermore, our engineered *E. coli* strain was designed to release HlyE toxin, with the intention of delivering the toxin directly to the site of the cancer cells and eliminating them. We successfully engineered *E. coli* HlyE Nb, a strain that displays Nb specific to HER2 receptors, which are commonly overexpressed in breast cancer cells, and that can secrete HlyE. We carried out a 2D assay that yielded favorable results, demonstrating the efficacy of our approach. To further corroborate and fortify our findings, we conducted a 3D assay that more closely mimics a tumor environment. Our 3D assay also showed promising results, providing evidence that our bacteria-based therapeutic approach is effective. The purpose of this inquiry was to devise a novel approach to deliver therapeutic agents with pinpoint precision to specific cells and tissues. It is worth noting that our methodology is adaptable to other cancer types, as nanobodies and toxin payloads can be customized for specific cancer cells. This adaptability offers a broad spectrum of potential therapeutic applications. Our study provides a compelling proof of concept for the use of bacteria-based therapeutics combined with nanotechnology for targeted cancer treatment. The potential for this approach is vast and holds tremendous promise for the development of new and effective cancer therapies. We envision that our work will pave the way for further research in this area, leading to more refined and sophisticated therapeutic strategies.

In terms of future prospects, the potential of bacteria-based therapy for cancer is tremendously encouraging, as ongoing research and development activities indicate that these inventive strategies could potentially transform cancer therapy. With the constant progress being made in the fields of synthetic biology, genetic engineering, biotechnology, and drug delivery systems, bacteria-based therapeutics offer a promising prospect of diversifying the available treatment options for cancer patients and enhancing the outcomes for those suffering from this disease. Bacteria possess the advantage of being readily modifiable in various ways and can be produced economically in bulk, making them an attractive candidate for therapeutic interventions. It is evident that the future of cancer therapy lies in the exploitation of bacteria, and the scientific community must continue to explore this avenue in order to translate these promising findings into practical solutions for patients.

## Materials and Methods

### Cloning and Plasmid Construction

The experiments for cloning utilized E. coli DH5α PRO cells. These cells were grown in Luria-Bertani (LB) medium, which consists of 10 g/l tryptone, 10 g/l NaCl, and 5 g/l yeast extract, with their corresponding antibiotics at 37 °C and 200 rpm. Overnight cultures were prepared from cell stocks stored in 25% (v/v) glycerol in LB medium and incubated in the same medium for 16 hours under the same conditions.

To construct the pET22b(+) T7-LacO PelB 2Rs15d Ag43 plasmid, a pET22b Ag43 160N 6h sfGFP AmpR plasmid was used as the template ^31^. The nanobody gene was amplified through Polymerase Chain Reaction (PCR) from the ordered gene fragment, followed by restriction digestion of the pET22b Ag43 160N sfGFP AmpR to obtain the backbone. The 2Rs15d nanobody was cloned downstream of the T7 LacO promoter using primers JO1 and JO2. After gel extraction, the parts were assembled through Gibson Assembly, and the final product was transformed into E. coli DH5α. Two colonies were selected for sequencing after verification of the correct plasmid. The plasmid was then transferred into E. coli BL21 (DE3) for the binding assay, which was double transformed with pZA mproD RFP for red fluorescence. The 2Rs15d nanobody was also cloned into an identical backbone containing a histidine tag to confirm its expression on the surface of E. coli BL21 (DE3) through Western Blotting (Figure S1).

To construct pTetO Yebf HlyE 6xHis, the backbone was taken from pZa HlyE YebF CmR vector, and HemolysinE was amplified from the same vector. The backbone was amplified using primers pREA29 and pREA78, and HemolysinE was amplified using JO29 and EK pBAD rev primers and a pZa pL-tetO YebF HlyE 6his CmR vector template. The parts were cloned through Gibson Assembly and chemically transformed into E. coli PRO DH5α. Colony PCR was performed to verify the cloning, and two colonies were selected for sequencing after validation.

To construct pZa native TetO Yebf HlyE 6xHis-mScarlet, the mScarlet insert was amplified from an Addgene plasmid using pGB1-pGB2 primers and cloned into the pTetO Yebf HlyE 6xHis backbone. The parts were assembled through Gibson Assembly and transformed into E. coli DH5α PRO. The cloning was verified through NGS sequencing. Double transformation was performed to obtain Bl21 pET22b T7 lacO his 2Rs15d Ag43 AmpR pZa native teto HlyE Yebf his CMR-mScarlet, followed by plasmid isolation and PCR. The mScarlet was also directly transformed into Bl21 competent cells to obtain Bl21-mScarlet. The sequences of the genetic parts and primers used in this study are provided in Tables S1 and S2. All of the constructs and maps for the cloning experiments and DNA sequence verificiation can be found in Figures S6 to S11.

### SDS-Page/Western Blotting

Samples were treated by boiling at 95°C for 5 minutes with 1x SDS loading buffer and separated by electrophoresis on a 15% SDS-polyacrylamide gel. The gel was stained with Coomassie blue solution for approximately 1 hour with continuous shaking and de-stained in a solution of 60% ddH2O, 30% methanol, and 10% acetic acid until the bands were visible. For Western blot analysis, the gel was transferred onto a PVDF membrane using a Trans-Blot Turbo apparatus (Bio-Rad). The membrane was then blocked for 1 hour at room temperature with 5% milk in TBS-T buffer. Next, the membrane was incubated overnight at 4°C with a 1:10,000 dilution of primary anti-His mouse antibodies in 5% milk-TBS-T. The membrane was washed with TBS-T and incubated for 1 hour at room temperature with a 1:10,000 dilution of HRP-conjugated goat anti-mouse secondary antibodies (Abcam ab6789-1 MG) in 5% milk-TBS-T. Followed by multiple washes by TBS-T, the membrane was treated with ECL substrates (Bio-Rad 170-5060) and visualized using a Vilber Fusion Solo S system.

Supernatant samples were prepared for Western blot analysis using trichloroacetic acid (TCA) precipitation. The cells were lysed in lysis buffer (50 mM Tris-HCl pH 7.5, 150 mM NaCl, 1% Triton X-100, 0.1% SDS, and 1 mM PMSF) for 30 minutes on ice. The lysates were then centrifuged at 14,000 g for 15 minutes at 4°C to remove insoluble debris. The resulting supernatants were mixed with 10% TCA and incubated at 4°C for 1 hour. The samples were then centrifuged at 14,000 g for 15 minutes at 4°C to pellet the precipitated proteins. The pellet was washed twice with ice-cold acetone and air-dried. The dried pellet was resuspended in loading buffer (0.125 M Tris-HCl pH 6.8, 4% SDS, 20% glycerol, and 0.2% bromophenol blue) and heated at 95°C for 5 minutes to denature the proteins. The samples were then separated by SDS-PAGE and transferred to a PVDF membrane for Western blot analysis.

### Immunocytochemistry

Antibodies were diluted 1:300 in a buffer containing 1% BSA in PBST (PBS + 0.1% Tween 20). After overnight incubation under inducing conditions, 5-10 mL of the cell culture was centrifuged at 4000 g for 15 minutes. The cell pellet was then treated with 1 mL of 4% formaldehyde diluted in PBS and incubated at room temperature for 30 minutes. For the washing step, the cells were centrifuged at 1500 g for 5 minutes and resuspended in 1 mL PBS two times. To prevent nonspecific binding of antibodies, slides were incubated in a blocking solution containing 1% BSA in PBST for 30 minutes. After removing the blocking solution, cells were incubated with the primary antibody for 2 hours at room temperature. Following washing in 500 μL of 1X PBS, the cells were incubated with a 1:500 dilution of secondary antibody overnight in the dark at +4 ºC. The cells were washed in PBS twice and left to air-dry. One drop of mounting medium was added to protect the staining and a cover glass was placed on each slide, fixed with nail polish. The slides were then visualized under a fluorescence microscope.

### Purification of HlyE protein

The HlyE protein was purified utilizing histidine tag columns, and the resulting eluted samples were analyzed through SDS-PAGE. The protein was found to be most concentrated in the second and third elution tubes. To make it suitable for use in mammalian cells, the protein was subjected to a buffer exchange with 1xPBS, as the initial buffer containing imidazole was toxic to these cells. After successful completion of the buffer exchange, the purified HlyE toxin was then tested on mammalian cells.

### Maintenance, Cultivation and Counting of JIMT1 cells

HER2+ breast cancer cell line JIMT1, stably transfected with GFP, was provided by the Molecular Biology and Genetics Department at Bilkent University, and stored at -80LC. Cells were cultivated in 88% low-glucose Dulbecco’s Modified Eagle Medium (DMEM), supplemented with 10% Fetal Bovine Serum (FBS), 1% Penicillin/ Streptomycin (Pen/Strep) antibiotic, and 1% L-glutamine. Cell stock was initially taken by adding contents of defrosted stock vial to 13 mL fresh media and centrifuged at 2500 rpm for 5 minutes. The supernatant was carefully discarded and the pellet was resuspended in 1 mL fresh medium. The suspension was added to a T25 flask together with 2 mL of media and incubated at 37°C, and 5% CO2. The media was changed every two days after washing with 3mL 1X PBS. Upon reaching 80%-90% confluency the cells where passaged. To passage the cells, they were first washed with 10 mL PBS, followed by addition of 3 mL trypsin, and incubating for 5 minutes to detach cells from flask surface. The contents were then added to 10 mL fresh medium in a 15 mL falcon and centrifuged for 5 minutes at 2500 rpm. The resulting supernatant was discarded and pellet dissolved in 1 mL fresh media. For cell counting, 10 μL of the sample was added into a PCR tube and diluted 1:10 with fresh media. 10 μL of this solution was added to both sides of a hemocytometer counting chamber covered by a glass coverslip, which was then viewed under inverted fluorescence microscopy at 20X magnification using bright-field illumination. The total number of cells in 1 mL was determined by counting the cells located inside the middle square (S1 and S2) bordered on four sides by three lines and applying the following formula: Total number of cells = 2 × (S1 + S2 + … + Snn) × 104 × dilution factor. To prepare the stock solution, one million cells were stored in 10% DMSO and stored overnight in an isopropanol chamber at - 80°C. The following day, the vial was transferred to the nitrogen tank for future use.

### Binding Assay

The localization of the bacteria to JIMT1 was determined using a binding assay. The pZA mproD RFP CmR plasmid, which mediates the expression of red fluorescence protein, was transformed into Bl21 pET22b T7 lacO his 2Rs15d Ag43 AmpR. The double-transformed bacteria were induced with 1 mM IPTG and centrifuged at 8000 rpm for 10 minutes, after which the media was discarded. The pellet was resuspended in 1X PBS under sterile conditions. *E. coli* Bl21 (DE3) were also inoculated as a control. JIMT1 cells cultured in a T75 flask were passaged, and 10^5 cells were added to each well of a 24-well plate in 100 µl of Low Glucose DMEM lacking FBS and antibiotics. A glass slide was placed at the bottom of each well. An equivalent number of CHO cells were added to wells in a separate well-plate as a control. 10^7 bacteria were counted and added to each well containing the mammalian cells, and the plates were incubated for one hour at 37°C. The glass slides were then removed, washed three times with 1X PBS to remove unbound bacteria, and placed on a drop of mounting solution on top of a microscope slide. Images were taken using a confocal microscope at 20X dry setting. The overall image showing the binding assay is given in Figure S2.

### 2D *in vitro* assay

JIMT1 cells grown in a T75 flask were passaged and counted as previously described. 10^4 cells were grown in 100 μL growth media and seeded in a 96 well plate to reach a confluency of 80% after overnight incubation at 37LC, and 5% CO2. Bacterial strains *E. coli* BL21 (DE3), pZa native teto HlyE Yebf his - mScarlet CmR, and Bl21 pET22b T7 lacO his 2Rs15d Ag43 AmpR pZa native teto HlyE Yebf his CMR – mScarlet were inoculated and induced with aTc and/or 1 mM IPTG. After induction, OD600 was measured for every sample in order to calculate the number of bacteria. The optical density at 600 nm (OD600) was measured for each sample to determine the number of bacteria, with an OD600 of 1 corresponding to approximately 10^9 bacteria (*E. coli*) per mL. The bacteria were then centrifuged at 8000 RPM 4°C for 10 minutes, and the supernatant was discarded. The bacterial pellet was suspended in sterile 1x PBS. JIMT1 wells were washed with 1X PBS and the media was replaced with 100 µL Low Glucose DMEM without Pen Strep and FBS. After making calculations, 5*10^5 bacteria cells were added to JIMT1 wells; the plate was incubated overnight at 37°C, 5% CO2. Confocal microscope pictures were taken the next day under 20X dry setting. Statistical analysis of the assay can be found in Figure S3.

### 3D cell culture

To assess the efficacy of our engineered bacteria in a 3D cell culture environment, we created JIMT1 cell spheroids using the agarose method. 1.5% High Purity Agarose was prepared by mixing 1.5 grams high purity agarose powder in 100 mL of PBS and sterilizing the mixture through autoclaving. 100 µl of the agarose solution was then added to wells in a 96-well plate and allowed to solidify for 1 hour. Next, JIMT1 cells were passaged and counted after being grown to 80% confluency in a T75 flask. Approximately 10,000-15,000 cells were added per well and the total volume was adjusted to 100µl by adding fresh media lacking Pen Strep and FBS. The cells were allowed to grow for 3 days to form into spheroids. The first set of images was taken before adding bacteria to the wells. The protocol for spheroid formation is shown in Figure S4.

The bacteria used in this experiment included *E. coli* BL21 - mScarlet, pZa native teto HlyE Yebf his - mScarlet CmR, and Bl21 pET22b T7 lacO his 2Rs15d Ag43 AmpR pZa native teto HlyE Yebf his CMR – mScarlet. The bacteria were inoculated, diluted and induced. After induction the bacterial suspensions were centrifuged for 5 minutes at 8000 rpm, and 4LC, and the supernatant was discarded while the pellet was resuspended in sterile 1X PBS. 10^5 bacterial cells were added to each well of the relevant groups. 0.3µl of aTc and IPTG were also added to the relevant wells. The plate was incubated at 37_C and the second set of images was taken after 4 hours. Imaging was done again over the next two days. All images were taken with Leica confocal microscope at 20X dry setting. And statistical analysis for the data can found in Figure S5.

### 3-(4,5-Dimethylthiazol-2-yl)-2,5-diphenyltetrazolium bromide (MTT) assay

3-(4,5-Dimethylthiazol-2-yl)-2,5-diphenyltetrazolium bromide (MTT) assay was performed to determine cell viability. The assay was performed in 96-well plates and each well was seeded with relevant number of cells per well in a final volume of 200 μL. After completion of experiment, the medium from wells was removed and replaced with 90 μL fresh medium plus 10 µL of MTT solution (5 mg/mL in PBS). The plates were then incubated for 4 hours at 37°C to allow for the reduction of MTT to formazan crystals. The resulting formazan crystals were solubilized in 100 μL of DMSO and the optical density was measured at 490/570 nm using a microplate reader.

## Statistical analysis

Statistical analysis was performed using GraphPad Prism 8.0 software. Multiple t-tests were conducted to compare the means of two groups. One-way ANOVA was performed to compare the means of multiple groups with a single factor, while two-way ANOVA was performed to compare the means of multiple groups with two factors.

