## Supplementary Material for "A Bacterial Living Therapeutics with Engineered Protein Secretion Circuits To Eliminate Breast Cancer Cells"

**Supplementary Figures :**

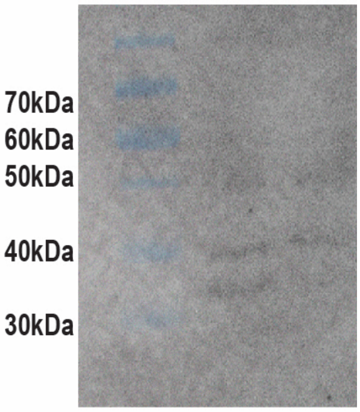


**Figure S1.** Western Blotting done to verify presence of 2Rs15d Nanobody (Nb) in engineered bacteria, *E. Coli* Nb. The relevant band is seen at ~40 kDa.

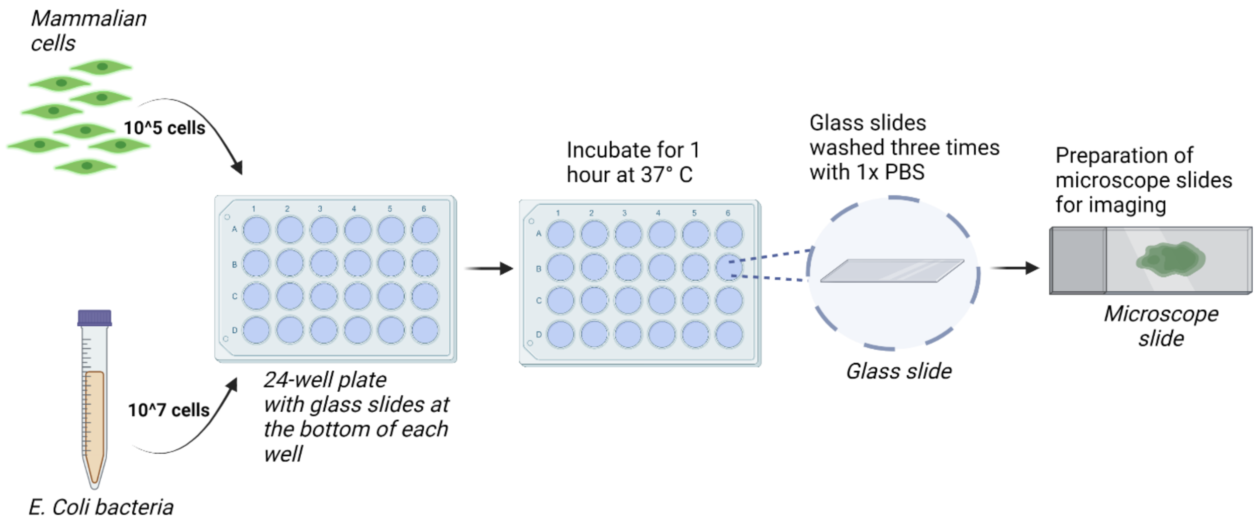


**Figure S2.** Workflow diagram for binding assay. Mammalian cells (JIMT1 and CHO) are seeded in a 24-well plate with each well containing a glass slide at the bottom. Bacteria *E. Coli* and *E. Coli* Nb is added to the wells after the mammalian cells have grown to 70% confluency. After 1 hour of incubation with bacteria, the glass slides are removed from the bottom of wells with the help of tweezers. These are washed with sterile 1X PBS three times to remove any unbound bacteria. The glass slides are then inverted and placed at the center of microscopic slides on a drop of mounting solution. They are secured with the help of nail varnish. The slides can then be stored at 4ºC or imaged.

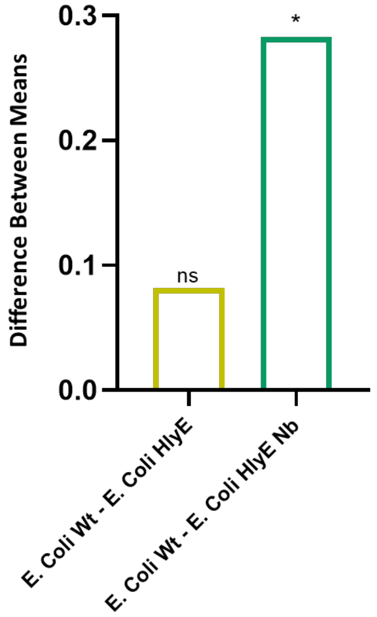

**Figure S3.** One-way Annova to compare 2D assay results between control *E. Coli* Wt and experimental *E. Coli* HlyE plus *E. Coli* HlyE Nb groups. We added Gentamycin to the wells containing bacteria and incubated the well-plate overnight before performing MTT assay. The cell viability readings were taken at absorbance value of 490 nm. The graph shows non-significant (ns) difference between means of 3 replicates of *E. Coli* Wt and *E. Coli* HlyE. As suspected there is a significant difference (*) between means of control group *E. Coli* Wt and *E. Coli* HlyE Nb. ($p > 0.05$ = ns, $p < 0.05$ = *).

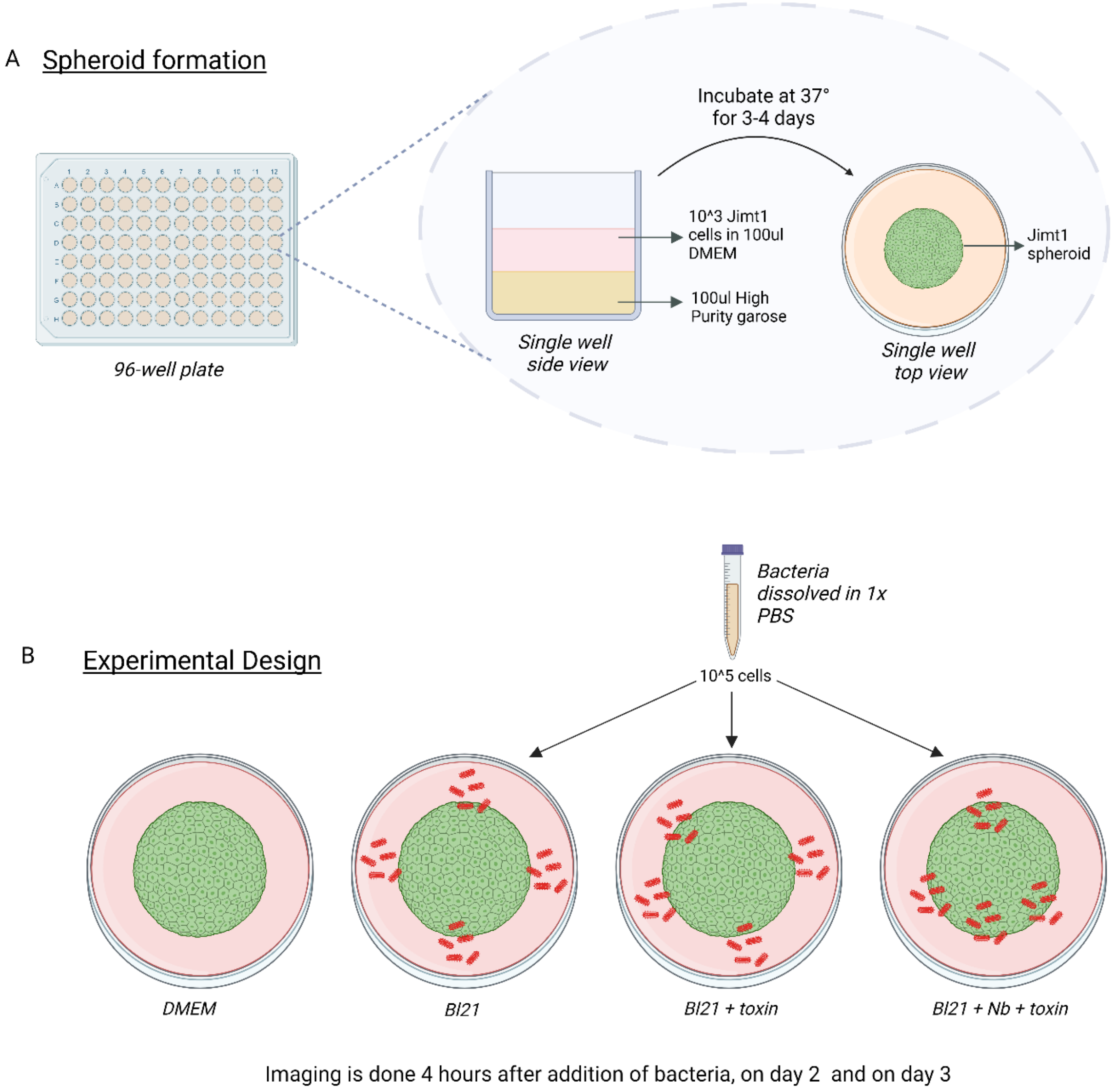


**Figure S4.** Spheroids experiment protocol. (A) Formation of spheroids using high purity agarose. 100 µl of high purity agarose was added to each well in a 96-well plate, and allowed to cool and solidify for one hour. Following this, approximately 10000-15000 JIMT1 cells were added per well and the total volume was made up to 100 µl with Low-Glucose DMEM without antibiotics. The cells were allowed to incubate for three days to form into spheroids. (B) Four replicate wells were made for each group: DMEM, *E. Coli* Wt, *E. Coli* HlyE and *E. Coli* HlyE Nb. Bacteria cultured in LB was induced with ATC and IPTG respectively. It was then centrifuged at 8000 rpm for 5 minutes at 4ºC. The supernatant was discarded and pellet resuspended in 1X sterile PBS. 10^5 bacterial cells were added to *E. Coli* Wt, *E. Coli* HlyE and *E. Coli* HlyE Nb groups ($OD_{600}$ of 1 = 10^9 cells). No bacteria was added in the DMEM group. Images were taken with Leica confocal microscope before addition of bacteria, 4 hours after addition of bacteria (Day 1), on Day 2, and on Day 3.

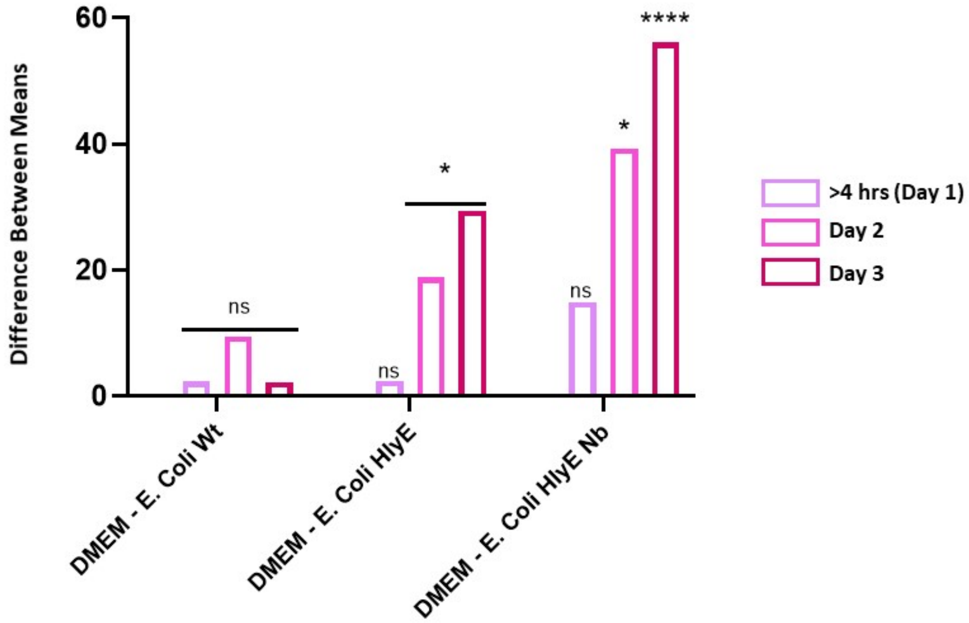


**Figure S5.** Two-way Annova using Dunnett's Muliple Comparisons Test ($p < 0.05$ = ns, $p < 0.0022$ = **, $p < 0.0001$ = ****, $p < 0.0001$ = ***), with DMEM as the control group for normalized spheroids areas over three days. DMEM and *E. Coli* Wt have a non-significant (ns) difference in mean spheroid areas over the three-day period. DMEM and *E. Coli* HlyE have a non-significant difference between means on day 1, however a significant difference (*) is reported on days 2 and 3. Compared to the control group DMEM, our main experimental group *E. Coli* HlyE Nb shows a significant difference in means of spheroid areas on day two (*) and a highly significant difference on day 3 (****), as hypothesized. This indicates that *E. Coli* HlyE Nb bacteria design is the most potent in disrupting JIMT1 cell spheroids.

**Supplementary Tables.**

**Table 1: Gene sequences used in this study**

| Name | Type | Sequence 5’-to-3’ |
|---|---|---|
| 2Rs15D | Nanobody | CAGGTGCAGCTGCAGGAAAGCGGCGGCGGCAGCGT<br>GCAGGCGGGCGGCAGCCTGAAACTGACCTGCGCGG<br>CGAGCGGCTATATTTTTAACAGCTGCGGCATGGGCT<br>GGTATCGCCAGAGCCCGGGCCGCGAACGCGAACTG<br>GTGAGCCGCATTAGCGGCGATGGCGATACCTGGCAT<br>AAAGAAAGCGTGAAAGGCCGCTTTACCATTAGCCAG<br>GATAACGTGAAAAAAACCCTGTATCTGCAGATGAACA<br>GCCTGAAACCGGAAGATACCGCGGTGTATTTTTGCG<br>CGGTGTGCTATAACCTGGAAACCTATTGGGGCCAGG<br>GCACCCAGGTGACCGTGAGCAGC |
| 2Rs15D ordered fragement | Nanobody | GCCCAGCCGGCGATGGCCATGGGCACTAGTCAGGT<br>GCAGCTGCAGGAAAGCGGCGGCGGCAGCGTGCAG<br>GCGGGCGGCAGCCTGAAACTGACCTGCGCGGCGAG<br>CGGCTATATTTTTAACAGCTGCGGCATGGGCTGGTA<br>TCGCCAGAGCCCGGGCCGCGAACGCGAACTGGTGA<br>GCCGCATTAGCGGCGATGGCGATACCTGGCATAAAG<br>AAAGCGTGAAAGGCCGCTTTACCATTAGCCAGGATA<br>ACGTGAAAAAAACCCTGTATCTGCAGATGAACAGCCT<br>GAAACCGGAAGATACCGCGGTGTATTTTTGCGCGGT<br>GTGCTATAACCTGGAAACCTATTGGGGCCAGGGCAC<br>CCAGGTGACCGTGAGCAGCCTTAAGCGCACAACCAT<br>CAATAAAAACGGT |
| pelB | Signal sequence | ATGAAATACCTGCTGCCGACCGCTGCTGCTGGTCTG<br>CTGCTCCTCGCTGCCCAGCCGGCGATGGCC |
| Ag43 (160 N alpha subunit) | Autotransporter Protein | CGCACAACCATCAATAAAAACGGTCGCCAGATTGTG<br>AGAGCTGAAGGAACGGCAAATACCACTGTGGTTTAT<br>GCCGGCGGCGACCAGACTGTACATGGTCACGCACT<br>GGATACCACGCTGAATGGGGGATACCAGTATGTGCA<br>CAACGGCGGTACAGCGTCTGACACTGTTGTGAACAG<br>TGACGGCTGGCAGATTGTCAAAAACGGGGGTGTGG<br>CCGGGAATACCACCGTTAATCAGAAGGGCAGACTGC<br>AGGTGGACGCCGGTGGTACAGCCACGAATGTCACC<br>CTGAAGCAGGGCGGCGCACTGGTTACCAGTACGGC<br>TGCAACCGTTACCGGCATAAACCGCCTGGGAGCATT<br>CTCTGTTGTGGAGGGTAAAGCTGATAATGTCGTACT<br>GGAAAATGGCGGACGCCTGGATGTGCTGACCGGAC<br>ACACAGCCACTAATACCCGCGTGGATGATGGCGGAA<br>CGCTGGATGTCCGCAACGGTGGCACCGCCACCACC<br>GTATCCATGGGAAATGGCGGTGTACTGCTGGCCGAT<br>TCCGGTGCCGCTGTCAGTGGTACCCGGAGCGACGG<br>AAAGGCATTCAGTATCGGAGGCGGTCAGGCGGATG<br>CCCTGATGCTGGAAAAAGGCAGTTCATTCACGCTGA<br>ACGCCGGTGATACGGCCACGGATACCACGGTAAATG<br>GCGGACTGTTCACCGCCAGGGGCGGCACACTGGCG |

| | | |
|---|---|---|
| | | GGCACCACCACGCTGAATAACGGCGCCATACTTACC CTTTCCGGGAAGACGGTGAACAACGATACCCTGACC ATCCGTGAAGGCGATGCACTCCTGCAGGGAGGCTCT CTCACCGGTAACGGCAGCGTGGAAAAATCAGGAAGT GGCACACTCACTGTCAGCAACACCACACTCACCCAG AAAGCCGTCAACCTGAATGAAGGCACGCTGACGCTG AACGACAGTACCGTCACCACGGATGTCATTGCTCAG CGCGGTACAGCCCTGAAGCTGACCGGCAGCACTGT GCTGAACGGTGCCATTGAC |
| Ag43 ( Beta subunit) | Autotransporter protein | CCCACGAATGTCACTCTCGCCTCCGGTGCCACCTGG AATATCCCCGATAACGCCACGGTGCAGTCGGTGGTG GATGACCTCAGCCATGCCGGACAGATTCATTTCACC TCCACCCGCACAGGGAAGTTCGTACCGGCAACCCTG AAAGTGAAAAACCTGAACGGACAGAATGGCACCATC AGCCTGCGTGTACGCCCGGATATGGCACAGAACAAT GCTGACAGACTGGTCATTGACGGCGGCAGGGCAAC CGGAAAAACCATCCTGAACCTGGTGAACGCCGGCAA CAGTGCGTCGGGGCTGGCGACCAGCGGTAAGGGTA TTCAGGTGGTGGAAGCCATTAACGGTGCCACCACGG AGGAAGGGGCCTTTGTCCAGGGGAACAGGCTGCAG GCCGGTGCCTTTAACTACTCCCTCAACCGGGACAGT GATGAGAGCTGGTATCTGCGCAGTGAAAATGCTTAT CGTGCAGAAGTCCCCCTGTATGCCTCCATGCTGACA CAGGCAATGGACTATGACCGGATTGTGGCAGGCTCC CGCAGCCATCAGACCGGTGTAAATGGTGAAAACAAC AGCGTCCGTCTCAGCATTCAGGGCGGTCATCTCGGT CACGATAACAATGGCGGTATTGCCCGTGGGGCCACG CCGGAAAGCAGCGGCAGCTATGGATTC GTCCGTCTGGAGGGTGACCTGATGAGAACAGAGGTT GCCGGTATGTCTGTGACCGCGGGGGTATATGGTGCT GCTGGCCATTCTTCCGTTGATGTTAAGGATGATGAC GGCTCCCGTGCCGGCACGGTCCGGGATGATGCCGG CAGCCTGGGCGGATACCTGAATCTGGTACACACGTC CTCCGGCCTGTGGGCTGACATTGTGGCACAGGGAA CCCGCCACAGCATGAAAGCGTCATCGGACAATAACG ACTTCCGCGCCCGGGGCTGGGGCTGGCTGGGCTCA CTGGAAACCGGTCTGCCCTTCAGTATCACTGACAAC CTGATGCTGGAGCCACAACTGCAGTATACCTGGCAG GGACTTTCCCTGGATGACGGTAAGGACAACGCCGGT TATGTGAAGTTCGGGCATGGCAGTGCACAACATGTG CGTGCCGGTTTCCGTCTGGGCAGCCACAACGATATG ACCTTTGGCGAAGGCACCTCATCCCGTGCCCCCCTG CGTGACAGTGCAAAACACAGTGTGAGTGAATTACCG GTGAACTGGTGGGTACAGCCTTCTGTTATCCGCACC TTCAGCTCCCGGGGAGATATGCGTGTGGGGACTTCC ACTGCAGGCAGCGGGATGACGTTCTCTCCCTCACAG AATGGCACATCACTGGACCTGCAGGCCGGACTGGAA GCCCGTGTCCGGGAAAATATCACCCTGGGCGTTCAG GCCGGTTATGCCCACAGCGTCAGCGGCAGCAGCGC TGAAGGGTATAACGGTCAGGCCACACTGAATGTGAC CTTC |

| | | |
|---|---|---|
| HlyE | Pore-forming Toxin | ATGACTGAAATCGTTGCAGATAAAACGGTAGAAGTAG TTAAAAACGCAATCGAAACCGCAGATGGAGCATTAG ATCTTTATAATAAATATCTCGATCAGGTCATCCCCTG GCAGACCTTTGATGAAACCATAAAAGAGTTAAGTCGC TTTAAACAGGAGTATTCACAGGCAGCCTCCGTTTTAG TCGGCGATATTAAAACCTTACTTATGGATAGCCAGGA TAAGTATTTTGAAGCAACCCAAACAGTGTATGAATGG TGTGGTGTTGCGACGCAATTGCTCGCAGCGTATATTT TGCTATTTGATGAGTACAATGAGAAGAAAGCATCCGC CCAGAAAGACATTCTCATTAAGGTACTGGATGACGG CATCACGAAGCTGAATGAAGCGCAAAAATCCCTGCT GGTAAGCTCACAAAGTTTCAACAACGCTTCCGGGAA ACTGCTGGCGTTAGATAGCCAGTTAACCAATGATTTT TCAGAAAAAAGCAGCTATTTCCAGTCACAGGTAGATA AAATCAGGAAGGAAGCATATGCCGGTGCCGCAGCC GGTGTCGTCGCCGGTCCATTTGGATTAATCATTTCCT ATTCTATTGCTGCGGGCGTAGTTGAAGGAAAACTGAT TCCAGAATTGAAGAACAAGTTAAAATCTGTGCAGAAT TTCTTTACCACCCTGTCTAACACGGTTAAACAAGCGA ATAAAGATATCGATGCCGCCAAATTGAAATTAACCAC CGAAATAGCCGCCATCGGTGAGATAAAAACGGAAAC TGAAACAACCAGATTCTACGTTGATTATGATGATTTAA TGCTTTCTTTGCTAAAAGAAGCGGCCAAAAAAATGAT TAACACCTGTAATGAGTATCAGAAAAGACACGGTAAA AAGACACTCTTTGAGGTACCTGAAGTC |
| YebF | Secretion system | ATGAAAAAAAGAGGGGCGTTTTTAGGGCTGTTGTTG GTTTCTGCCTGCGCATCAGTTTTCGCTGCCAATAATG AAACCAGCAAGTCGGTCACTTTCCCAAAGTGTGAAG ATCTGGATGCTGCCGGAATTGCCGCGAGCGTAAAAC GTGATTATCAACAAAATCGCGTGGCGCGTTGGGCAG ATGATCAAAAAATTGTCGGTCAGGCCGATCCCGTGG CTTGGGTCAGTTTGCAGGACATTCAGGGTAAAGATG ATAAATGGTCAGTACCGCTAACCGTGCGTGGTAAAA GTGCCGATATTCATTACCAGGTCAGCGTGGACTGCA AAGCGGGAATGGCGGAATATCAGCGGCGT |
| Sec | Signal peptide | ATGAAAAAAAGAGGGGCGTTTTTAGGGCTGTTGTTG GTTTCTGCCTGCGCATCAGTTTTCGCT |
| T7 promoter | Promoter | TAATACGACTCACTATAGG |
| LacO | Promoter | GGAATTGTGAGCGGATAACAATTCC |
| T7 terminator | Terminator | CTAGCATAACCCCTTGGGGCCTCTAAACGGGTCTTG AGGGGTTTTTTG |
| rrnB T1 | Terminator | CAAATAAAACGAAAGGCTCAGTCGAAAGACTGGGCC TTTCGTTTTATCTGTTGTTTGTCGGTGAACGCTCTCCT GAGTAGGACAAAT |
| PL-TetO | Promoter | TCCCTATCAGTGATAGAGATTGACATCCCTATCAGTG ATAGAGATACTGAGCACATCAGCAGGACGCACTGAC C |

| | | |
|---|---|---|
| GST tag | Protein | ATGTCCCCTATACTAGGTTATTGGAAAATTAAGGGCCTTGTGCAACCCACTCGACTTCTTTTGGAATATCTTGAAGAAAAATATGAAGAGCATTTGTATGAGCGCGATGAAGGTGATAAATGGCGAAACAAAAAGTTTGAATTGGGTTTGGAGTTTCCCAATCTTCCTTATTATATTGATGGTGATGTTAAATTAACACAGTCTATGGCCATCATACGTTATATAGCTGACAAGCACAACATGTTGGGTGGTTGTCCAAAAGAGCGTGCAGAGATTTCAATGCTTGAAGGAGCGGTTTTGGATATTAGATACGGTGTTTCGAGAATTGCATATAGTAAAGACTTTGAAACTCTCAAAGTTGATTTTCTTAGCAAGCTACCTGAAATGCTGAAAATGTTCGAAGATCGTTTATGTCATAAAACATATTTAAATGGTGATCATGTAACCCATCCTGACTTCATGTTGTATGACGCTCTTGATGTTGTTTTATACATGGACCCAATGTGCCTGGATGCGTTCCCAAAATTAGTTTGTTTTAAAAAACGTATTGAAGCTATCCCACAAATTGATAAGTACTTGAAATCCAGCAAGTATATAGCATGGCCTTTGCAGGGCTGGCAAGCCACGTTTGGTGGTGGCGACCATCCTCCAAAA |
| mScarlet | Fluorescent protein | ATGAGTAAAGGAGAAGCTGTGATTAAAGAGTTCATGCGCTTCAAAGTTCACATGGAGGGTTCTATGAACGGTCACGAGTTCGAGATCGAAGGCGAAGGCGAGGGCCGTCCGTATGAAGGCACCCAGACCGCCAAACTGAAAGTGACTAAAGGCGGCCCGCTGCCTTTTTCCTGGGACATCCTGAGCCCGCAATTTATGTACGGTTCTAGGGCGTTCACCAAACACCCAGCGGATATCCCGGACTATTATAAGCAGTCTTTTCCGGAAGGTTTCAAGTGGGAACGCGTAATGAATTTTGAAGATGGTGGTGCCGTGACCGTCACTCAGGACACCTCCCTGGAGGATGGCACCCTGATCTATAAAGTTAAACTGCGTGGTACTAATTTTCCACCTGATGGCCCGGTGATGCAGAAAAAGACGATGGGTTGGGAGGCGTCTACCGAACGCTTGTATCCGGAAGATGGTGTGCTGAAAGGCGACATTAAAATGGCCCTGCGCCTGAAAGATGGCGGCCGCTATCTGGCTGACTTCAAAACCACGTACAAAGCCAAGAAACCTGTGCAGATGCCTGGCGCGTACAATGTGGACCGCAAACTGGACATCACCTCTCATAATGAAGATTATACGGTGGTAGAGCAATATGAGCGCTCCGAGGGTCGTCATTCTACCGGTGGCATGGATGAACTATACAAATAA |

**Table S2.** List of primers used in this study

| Name | DNA Sequence (5'-to-3') |
|---|---|
| JO1 | ACCGTTTTTATTGATGGTTGTGCGCTTAAGGCTGCTCACGGTCACCTG |
| JO2 | CACCACCACCACCACGGTGGCGGTGGATCCCAGGTGCAGCTGCAGGAAAG |
| JO5 | GGTGGTGGCGACCATCCTCCAAAAGGATCCATGACTGAAATCGTTGCAGATAAAAC |
| JO6 | TCTCAGTGGTGGTGGTGGTGGTGCTCGAGTCATCAGACTTCAGGTACCTCAAAGAG |
| PREA24 | TCTGTTCCAGGGGCCCCTGGGATCCGGTGGCGGTGGTAGCGAAAACCTGTACTTTCAGGGCGGTG |
| MKL2 | CCGTTTTTATTGATGGTTGTGCGCTTAAGATTCATCAGCTGGGTGGCTT |
| pZA Ag43 atc FWD | CTTAAGCGCACAACCATCAATAAAAACGGTCG |
| pZA Ag43 atc REV | CACGGTGGCGGTGGATCC |
| JO11 | CGCTGCTGCTGGTCTGCTGCTCCTCGCTGCCCAGCCGGCGATGGCCGGTGGCGGTGGATCCATG |
| JO12 | TTCGTTTTATTTGATGCCACGCGTCTCGAGTCATTAGTGGTGGTGGTGGTGGT |
| PREA29 | ACGCGTGGCATCAAATAAAACGAAA |
| PREA78 | CTTTTTTTCATGGTACCCTCCTTCAGATCTTTTGAATTCTTTTCTCTATC |
| JO29 | AAGAATTCAAAAGATCTGAAGGAGGGTACCATGAAAAAAAGAGGGGCGTT |
| pGB1 | CTTTTTTATTATTTCCTGCAGGGCGGCCGCCCACAAGCTAACACCACGTCGTC |
| pGB2 | TCGAGATTTTCAGGAGCTAAGGAAGCTAAAGCATAAAACGAAAGGCCCAGTCTTTCGA |

### DNA Sequences Verification of the Genetic Parts Used in the Study

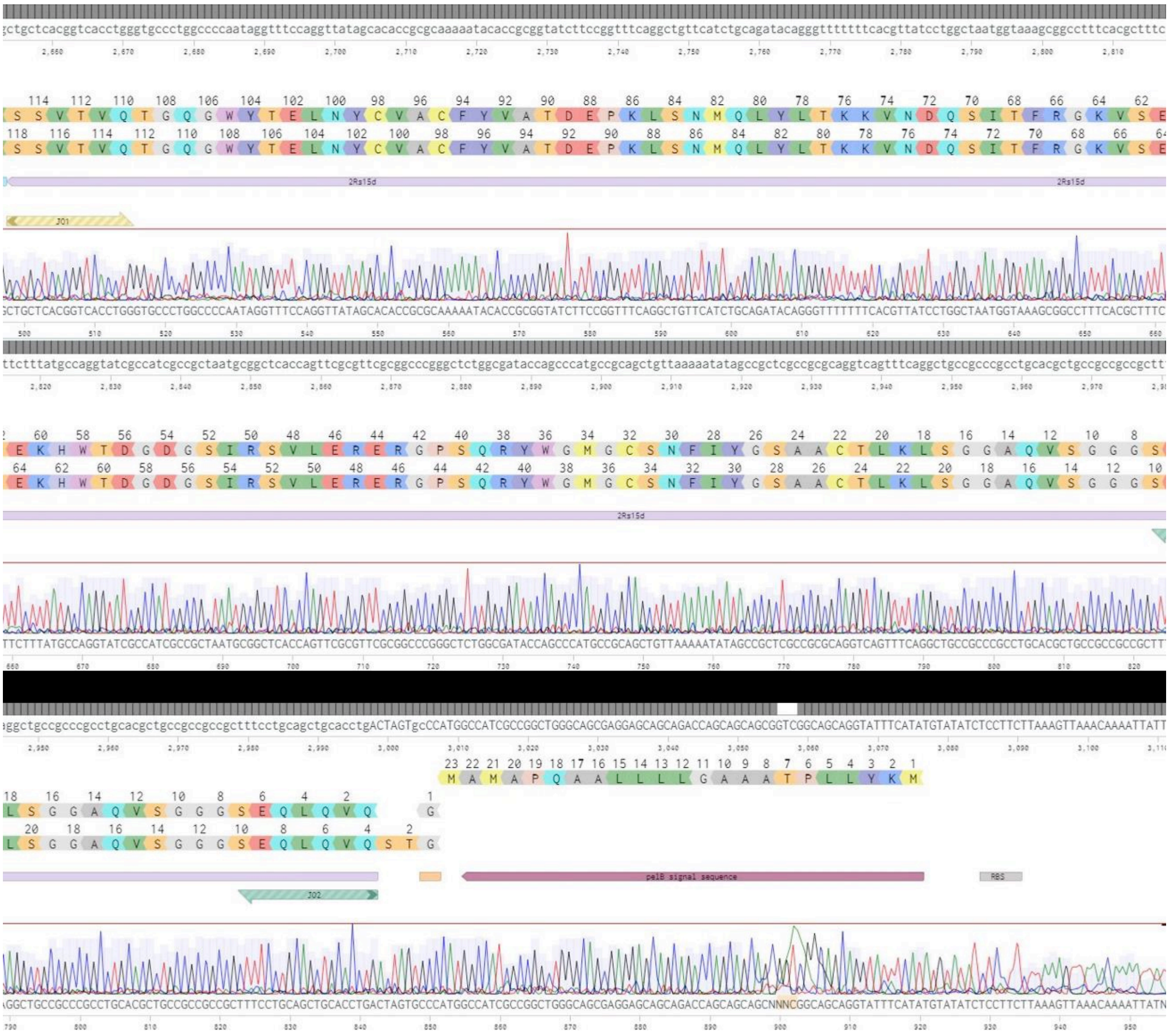

**Figure S6.** Alignment of pET22b T7-LacO pelB 2Rs15d Ag43 AmpR with sanger sequencing result via Benchling.

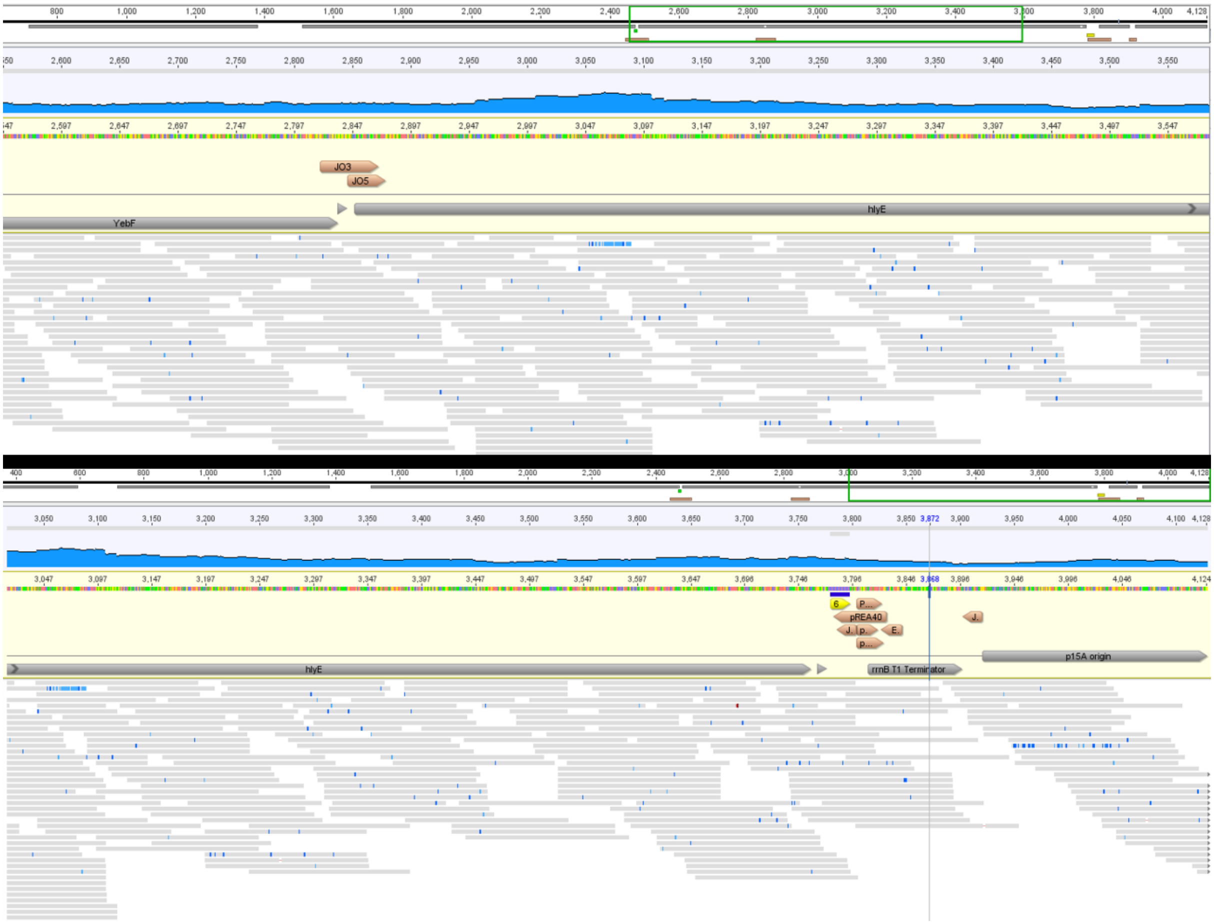


**Figure S7.** Alignment of pZA native pTetO HlyE Yebf 6xHis CmR sequencing result using Geneious program.

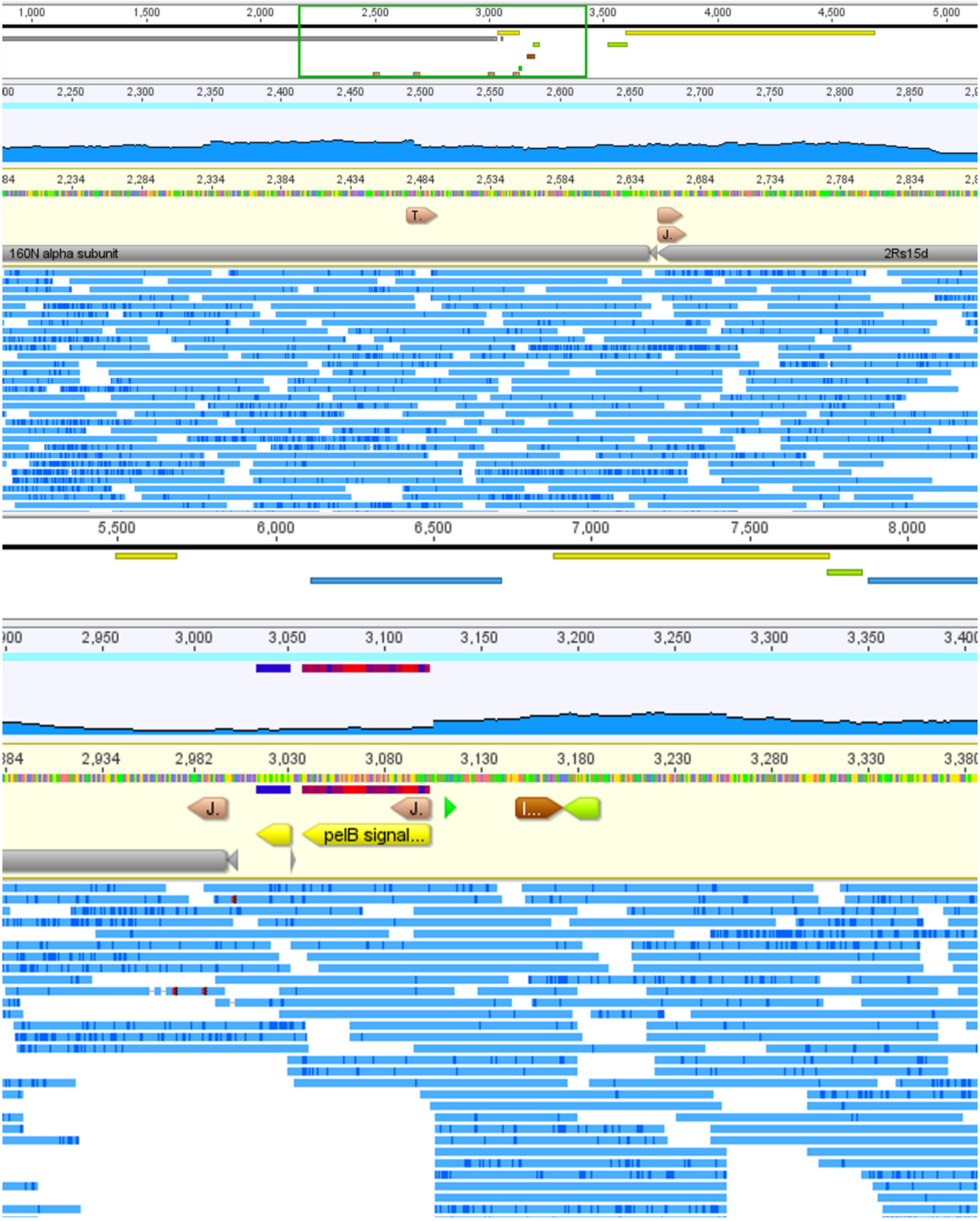


**Figure S8.** Alignment of pET22b T7-LacO pelB 6xHis 2Rs15d Ag43 AmpR sequencing result using Geneious program.

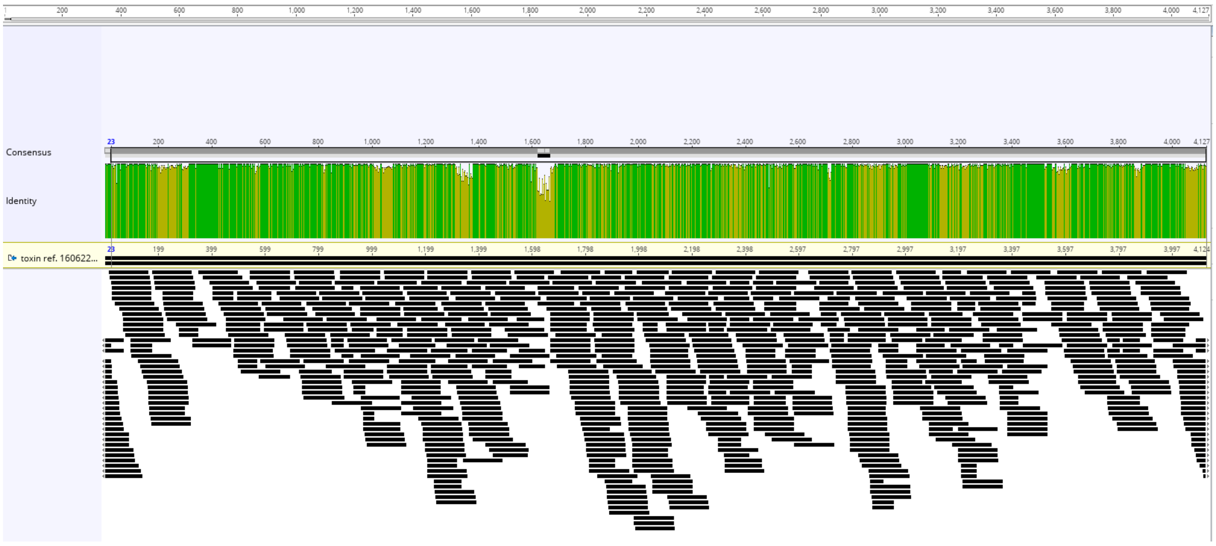

**Figure S9.** Alignment of pZA native pTetO HlyE Yebf 6xHis CmR – mScarlet using NGS sequencing.

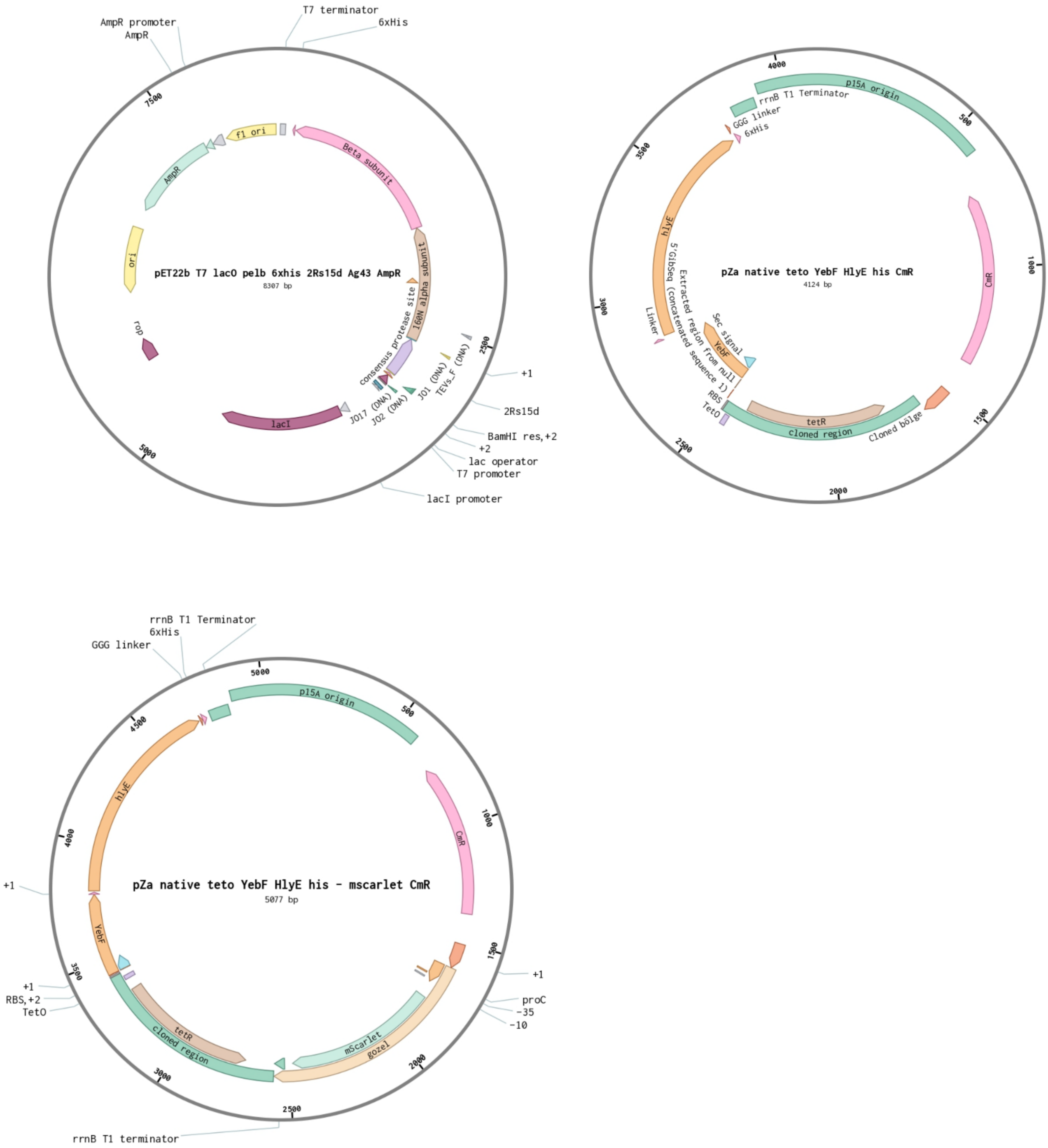


**Figure S10.** Plasmids designed on Benchling.com used in this study.
